# A systematic investigation of human DNA preservation in medieval skeletons

**DOI:** 10.1101/2020.05.20.106971

**Authors:** Cody Parker, Adam B. Rohrlach, Susanne Friederich, Sarah Nagel, Matthias Meyer, Johannes Krause, Kirsten I. Bos, Wolfgang Haak

## Abstract

Ancient DNA (aDNA) analyses necessitate the destructive sampling of archaeological material. Currently the dense inner portion of the petrous pyramid, the location of the skull that houses the inner ear, is the most sought after skeletal element for molecular analyses of ancient humans as it has been shown to yield high amounts of endogenous DNA. Destructive sampling of the petrous pyramid, assuming its recovery, is often not recommended for highly valued specimens. To investigate alternatives, we present a survey of human aDNA preservation for each of ten skeletal elements in a skeletal collection from Medieval Germany. Through comparison of human DNA content and quality we confirm best performance of the petrous pyramid and identify seven additional sampling locations across four skeletal elements that yield adequate aDNA for most applications in human palaeogenetics. Our study provides a better perspective on DNA preservation across the human skeleton and takes a further step toward the more responsible use of ancient materials in human aDNA studies.

## Introduction

The study of ancient DNA (aDNA) has progressed rapidly over the past decade following the introduction of next generation sequencing^1–3^, where genome-level analyses of archaeological specimens are now standard^4–12^. The increased analytical resolution offered by large scale datasets, coupled with the establishment of laboratory techniques that permit parallel processing of large sample sizes, has resulted in an increasing demand for ancient skeletal samples for assessment of human population genetics, microbiome ecology, and investigations of pathogen evolution. Laboratory processing of ancient remains is intrinsically a destructive process^13–16^, which poses ethical challenges related to the use of irreplaceable resources. Coupled with the high processing costs of aDNA work (from the perspective of both financial and time investments), there is benefit in optimizing approaches for material sampling. Multiple investigations have demonstrated superior human aDNA preservation in the dense inner petrous pyramid, the portion of the temporal bone that houses the inner ear. This observation is based on a collection of comparative PCR^15,17–20^ and whole genome aDNA surveys^16–18^ that were, however, limited in the number of individuals and/or skeletal elements tested. Despite the absence of a systematic comparative analysis of preservation across the skeleton, aDNA obtained from the petrous portions of human remains has been utilized to great success in the contexts of both ancient human population genetics^21–25^ and forensic investigations^12,24,25^.

Historically, sampling of the isolated petrous pyramid has typically involved sectioning or sand-blasting of the temporal bone to isolate the cochlea^16^, making this a highly destructive process^13^. Recent advances in minimally invasive sampling techniques^26^ have led to a better balance between preservation of the anthropological record and the need for the production of reliable genetic data^27,28^; however, the threat of damage to internal microstructures that form an important basis of morphological assessments^29–31^ can still introduce hesitancy on the part of curators and physical anthropologists in making the petrous pyramid available for aDNA applications. These factors, in conjunction with the chance of incomplete recovery of crania at excavation^32^ or restricted sampling of highly valued specimens, make the identification of alternative sampling locations based on quantitative evaluations of DNA preservation across the skeleton of clear benefit. Teeth have been widely used for the study of aDNA^33,34^, though the 30-fold covered genome of an archaic hominin from Denisova Cave from a distal phalanx demonstrates molecular preservation in elements that are not typically considered for paleogenetics work^4^. Despite these successes, a systematic and extensive study of differential DNA preservation across multiple human skeletal elements, such as those done in the context of modern forensics^35,36^, has yet to be attempted on archaeological remains. Our limited understanding of DNA preservation across the human skeleton is a significant hurdle for the efficient, practical, and ethical study of aDNA, which has particular relevance to the field of ancient population genetics where large sample sizes are needed for robust analytical resolution.

DNA preservation can be influenced by many factors including burial practises and treatment of the deceased, geology, as well as environmental and climatic conditions^37^, where the chronological age of a sample is thought to play only a secondary role^38,39^. To serve as a baseline for future investigations seeking to incorporate and extrapolate the effects of these sources of variation, e.g. across other species, time series, or geographic regions we present a broad survey of aDNA preservation across a long list of skeletal elements. Our source material, consequently, has been deliberately restricted to one archaeological site and time period to control for these factors that can influence molecular recovery as much as possible. The range of elements chosen for this survey consist of petrous bones (chosen for their demonstrated value in aDNA recovery^40,41^), *in situ* molars, clavicles, the first ribs, thoracic vertebrae, metacarpals, distal phalanges, ischial tuberosities, femora (once widely used in ancient DNA studies^42^), and tali. Multiple locations on each element were drilled and evaluated for DNA content. A detailed list of skeletal elements, sampling locations and the rationale for why each element was selected for study is provided in Table 1 (see also Supplementary Material: Section 1.2). Differential DNA preservation across these elements was investigated in individuals excavated from the church cemetery associated with the abandoned medieval settlement of Krakauer Berg, near Peißen, Saxony-Anhalt, Germany (Figure 1). Overall, the site exhibited high levels of morphological preservation, and as such, sampling from complete (or nearly complete) skeletons permitted us to maximize the comparability of the elements selected for analyses while also maximizing the chances of successful DNA extraction from each sampling location, thus minimizing the number of potential missing data points.

**Table 1.**
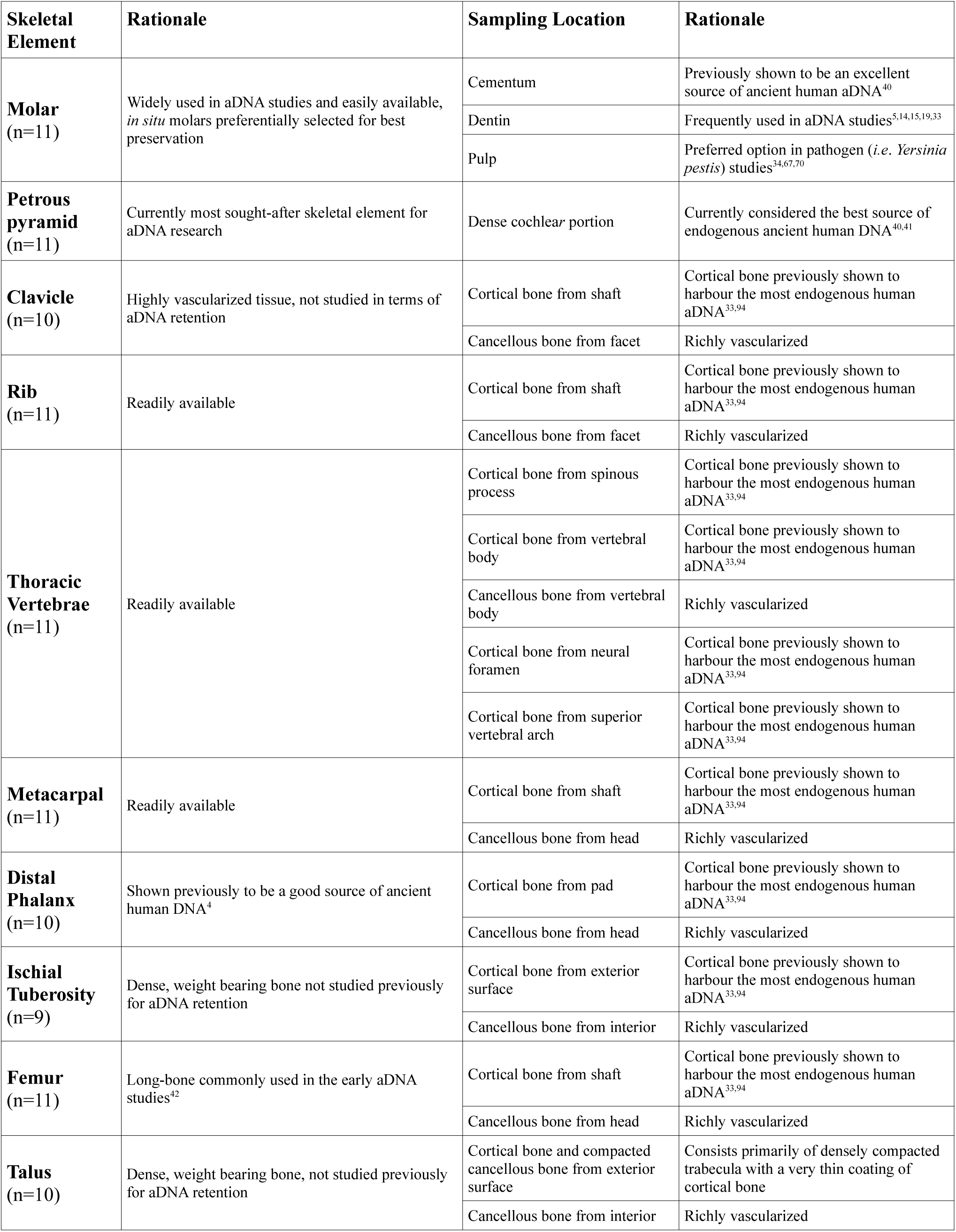
Skeletal elements, sampling locations as well as the rationale behind the choice of element.

**Figure 1.**
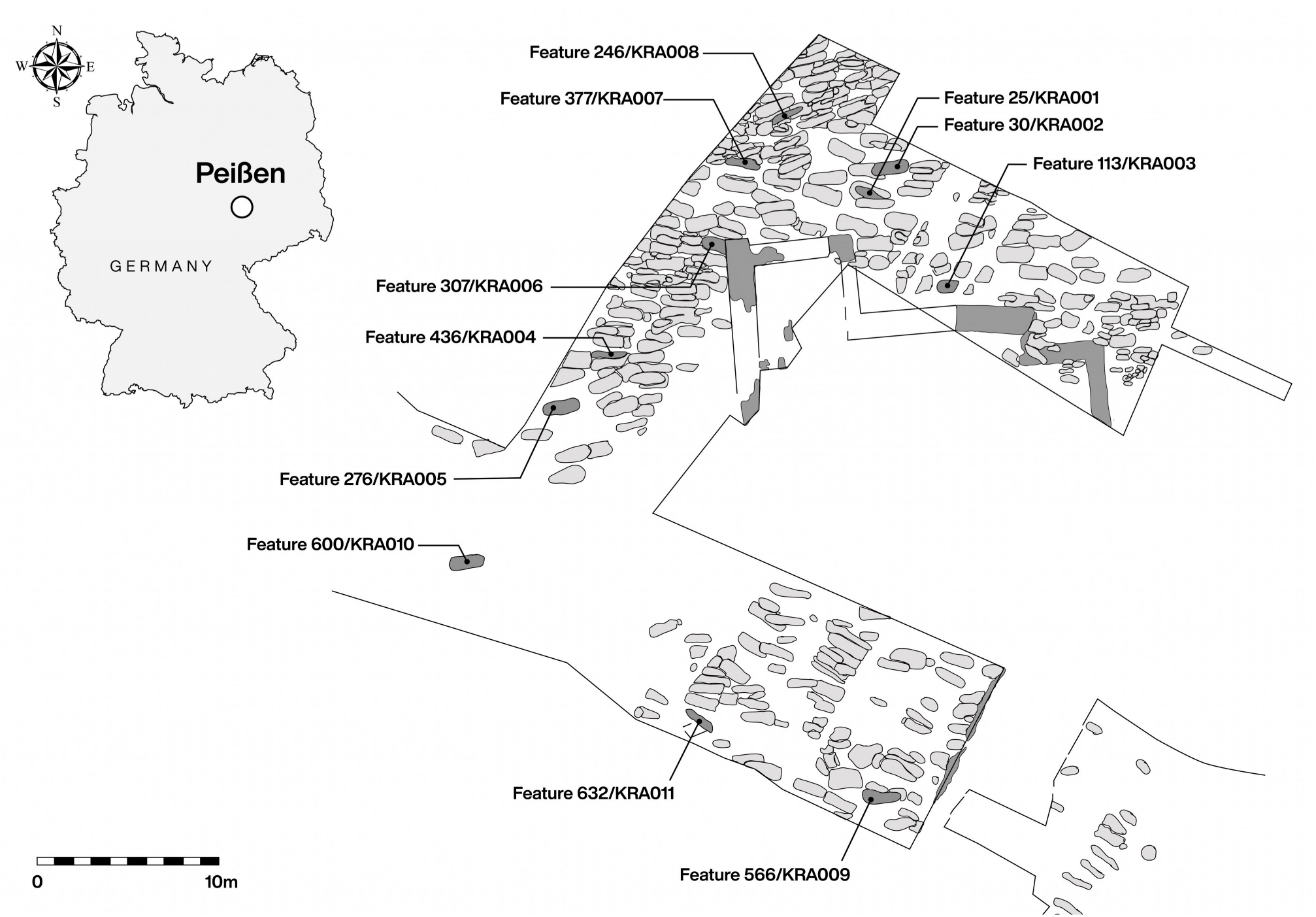
Map of the Krakauer Berg excavation. Graves corresponding to individuals sampled are denoted with both the archaeological ID and assigned sample name.

To our knowledge, this study presents the most comprehensive evaluation of aDNA preservation across the human skeleton in the published literature. While we further confirm the superior performance of cortical bone stemming from the cochlear portion of the petrous pyramid to yield the highest amounts of recoverable human DNA^40,41,43^, several alternative sampling locations are identified as suitable for downstream population genetic analyses such as the tali, distal phalanges, vertebrae, and teeth.

## Results

Our analytical matrix consists of shotgun sequencing data (ca. 5,000,000 reads at 75bp, paired end for each single-stranded library) from 23 separate sampling locations (Table 1; Supplementary File 1: Source; Raw reads sequenced). These were obtained from ten skeletal elements from each of eleven individuals who were all buried, excavated, documented, stored, sampled from, and ultimately processed and sequenced under the same conditions, in order to eliminate as many confounding variables as possible. All individuals selected for study had at least nine elements available, and all elements were present in at least nine individuals. In total, this resulted in 246 single-stranded aDNA libraries for comparison. In addition, as the use of hybridization capture technology is gaining popularity as a cost-effective and accessible alternative to shotgun sequencing^25^ an additional 87 libraries were subsequently enriched by hybridization capture for 1,240,000 informative variant SNPs across the human genome using the 1240k^25^ human SNP array and sequenced to a depth of ca. 40 million reads each (75bp, paired end). Our goals in evaluating this dataset are to ascertain which of the chosen sampling locations are most efficient in terms of authentic host DNA recovery, processing cost, and limiting damage to the anthropological record. To achieve a balance of data and drilling damage, as well as to more accurately compare the expected yields from a single instance of sampling, each sampling location was screened only one time on each bone (analyses normalized in terms of input material available from each sampling locations can be found in the provided Supplementary text in Section 2.4).

One of the most frequently used metrics for the evaluation of successful DNA recovery in human archaeological material is the proportion of human DNA recovered relative to DNA from other sources. This is often the first criterion considered to determine if a sample is suitable (both economically and analytically) for further testing. In this context we examined the average proportion of total (prior to duplicate removal) human DNA recovered post paired-end read merging, accommodating filters for sequence length and mapping quality (see Methods: Calculations). Among the 23 sampling locations we find the highest average proportion of human DNA in the petrous pyramid (34.70% human DNA on average), followed by dense tissue obtained from the neck and articular surfaces of the talus (21.25%), the cementum (18.97%), cortical bone from the distal phalanx (18.89%), material from the dental pulp chamber (15.09%), cortical bone from the vertebral body (15.04%), the dentin (14.27%), and cortical bone from the superior vertebral arch (8.32%). All other sampling locations evaluated contained an average human DNA proportion lower than the overall average of 8.16% (Figure 2A, Supplementary File 1: % mapping q37) across all elements tested.

**Figure 2A-C.**
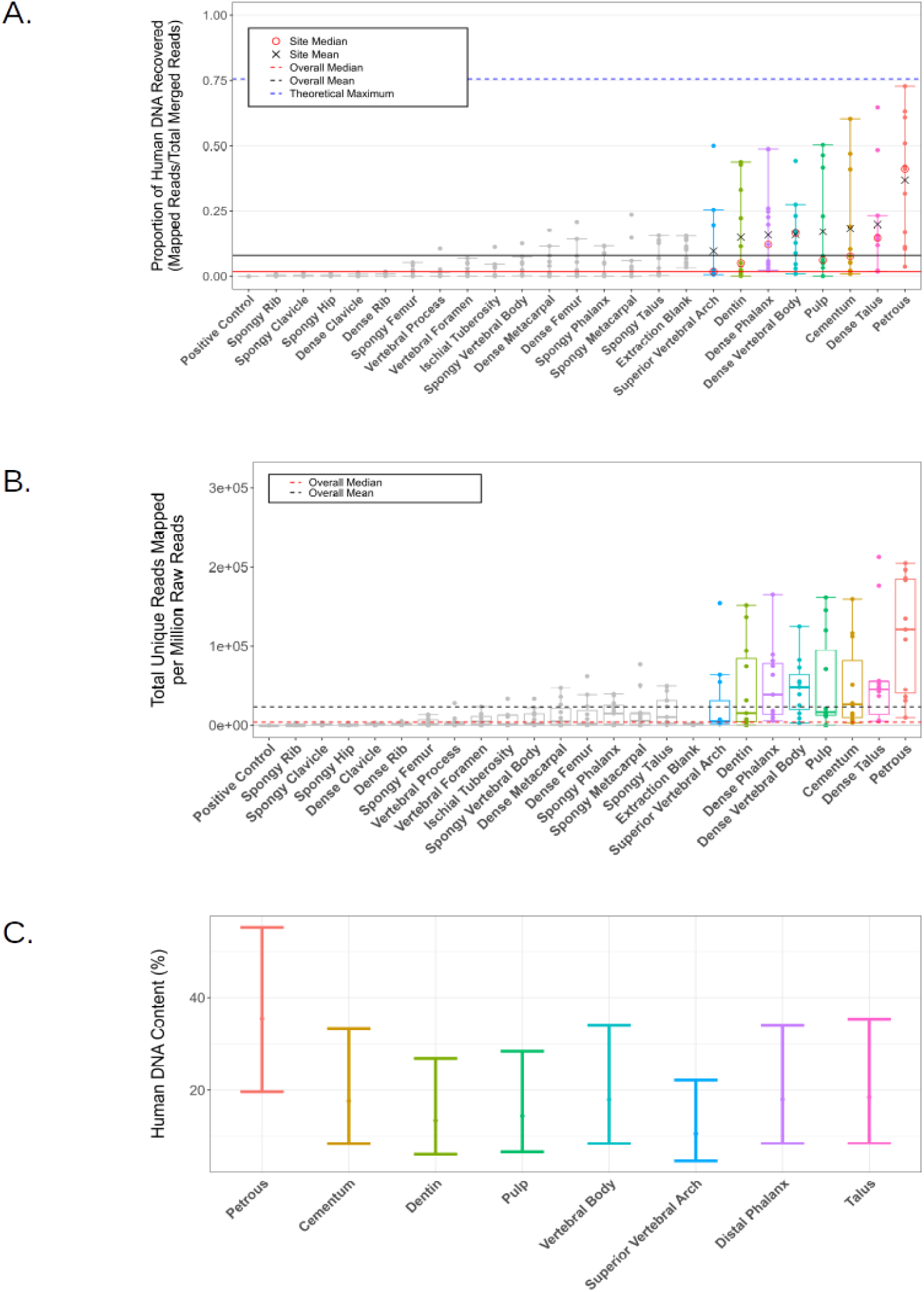
Human DNA content for all screened samples. Black lines represent the overall mean, red the median (solid: human DNA proportion, dashed: mapped human reads per million reads generated). Individual sampling locations with an average human DNA proportion higher than the overall mean (8.16%) are colourized in all analyses. **A)** Proportion of reads mapping to the hg19 reference genome. The blue dashed line represents the theoretical maximum given the pipeline’s mapping parameters (generated using Gargammel^89^ to simulate a random distribution of 5,000,000 reads from the hg19 reference genome with simulated damage). Individual means (black X) and medians (red circle) are reported for those samples with a higher average human DNA proportion than the overall mean. **B)** Number of unique reads mapping to the hg19 reference genome per million reads of sequencing effort (75bp paired end Illumina). **C)** Predicted range of expected human DNA recovery (in proportion of total reads) for each top scoring sampling site. Predictions were generated using a beta-fitted mixed effects model to simulate 55,0000 sampling iterations.

To provide a realistic approximation of the cost efficiency of human DNA retrieval from each sampling location, we further compared the average number of unique human reads per million reads of sequencing effort across all samples (see Methods: Calculations). Here we again find the highest average in the petrous pyramid (1.14×10^5^ unique reads mapping per million), followed by the talus (6.43×10^4^ unique reads mapping per million), the dental pulp chamber (5.26×10^4^ unique reads mapping per million), the distal phalanx (5.23×10^4^ unique reads mapping per million), cementum (4.89×10^4^ unique reads mapping per million), the vertebral body (4.81×10^4^ unique reads mapping per million), the dentin (4.76×10^4^ unique reads mapping per million), and the superior vertebral arch (2.79×10^4^ unique reads mapping per million). All other sampling locations fall below the overall average of 2.43×10^4^ (Figure 2B, Supplementary File 1: Unique reads/million reads). Among these sampling locations an average unique reads/million lower than that found in the highest of our extraction blanks (2.96×10^3^ unqiue reads per million) was observed in all sampling locations on the ribs and clavicula, as well as cancellous material from the ischial tuberosities. When normalized to reflect the amount of input material from each sampling effort, we find those sampling locations with the lowest available input material to yield the highest average number of unique mapping reads per million per mg of input material, followed by the petrous pyramid (cementum: 3751 unique reads mapping/million reads/mg, material from the pulp chamber: 2736 unique reads mapping/million reads/mg, and petrous pyramid 2087 unique reads mapping/million reads/mg) (Supplementary Figure S14, Supplementary File 1: Unique reads/mg/million), suggesting that material from the cementum and dental pulp chamber may be especially rich in human DNA.

It should also be noted that, while human DNA content in the negative controls was relatively high on average (10.77%), this metric is not directly informative for the evaluation of potential contamination as there are comparatively few DNA molecules in negative controls and as a result high numbers of amplification rounds are typically required, yielding an abundance of clonal PCR duplicates (see Supplementary File 1: Reads raw sequencing effort, Reads after merging, and Unique reads/million reads). The number of unique mapping reads per million is, therefore, a more informative metric. Here the average among our controls is an order of magnitude lower than what we report for our samples (an average of 1,67×10^3^ unique reads mapping per million in extraction blanks vs an average of 2.43×10^4^ unique reads mapping per million reads overall; see Supplementary File 1: Unique reads/million reads). Using a conservative approach, we considered all individual sampling efforts that yielded a lower number of unique reads/million than what was observed in the highest of the negative controls (2.96×10^3^ unique reads mapping/million reads) to be unsuccessful, regardless of potential authenticity as determined by characteristic patterns of DNA decay typically indicating ancient origin. (see Supplementary File 1: Damage signals). With this in mind, however, all “failing” samples were retained for all downstream comparative analyses so as to more accurately represent the expected outcomes of sampling efforts across a given sampling location. We additionally observed that all cancellous samples, as well as cortical bone samples stemming from ribs, claviculae, metacarpals, ischial tuberosities, femora, neural foramen and spinous process of the thoracic vertebrae (15 sampling locations, n=158) exhibited average human DNA contents lower than the overall averages (>8.16% for human DNA proportion, and 2.43×10^4^ for unique human reads/million reads) making them unlikely to be among the most efficient sampling locations in any metric. Accordingly, we removed these sampling locations from further analyses to allow for the deeper investigation of the remaining eight sampling locations consisting of the dentin, cementum, and dental pulp chambers as well as cortical bone from the cochlear portion of the petrous pyramid, vertebral body, superior vertebral arch, distal phalanx, and talus (eight sampling locations, n=87).

Restriction of our dataset to these eight sampling locations also permitted generation of a predictive model of expected human DNA yields via mixed effects beta regression (Figure 2C). Using this approach, we were able to take into account unavoidable sources of variation such as those stemming from individual preservation at particular skeletal locations (*i.e*. the natural variability among sampling locations across individuals). Due to the high variability of the proportion of human DNA recovered across both sampling locations and individuals, 55,000 iterations of this simulation were run to evaluate overall consistency of the expected proportion of human DNA recovered from each sampling location (Supplemental Material: Table S1). Here, the petrous pyramid significantly outperformed all other tested elements in terms of the expected range of proportions of recovered human DNA (all p-values < 0.0279), and yielded the highest predicted proportion of human DNA in the greatest number of simulations (41.87% of 55,000 simulations). The seven remaining alternative sampling locations on four other elements, although second to the petrous pyramid, also exhibit excellent human DNA recovery with yields statistically indistinguishable from each other (p-values > 0.1) (Figure 2C). The distal phalanx, vertebral body, cementum and talus yielded the highest proportion of human DNA in 9.93-10.61% of simulations, followed by the pulp chamber, dentin, and superior vertebral arch, which yielded the highest proportions in 4.28-7.22% of the simulations.

Although the proportion of human DNA is vitally important for the identification of suitable sampling locations, both the quantity and quality of that DNA are also important for the success of downstream analyses. With that in mind, we examined several additional aspects of DNA preservation. As many studies require robust assignments of genetic variants at individual loci, it is important that aDNA libraries are of sufficient complexity and show low signals of contamination with present-day human DNA. The aDNA libraries produced in this study were not sequenced to exhaustion, and as a consequence duplication rates were too low to be informative in terms of estimating library complexity in both the pre-enrichment libraries (average duplication factor 1.21) and the post-capture libraries (average duplication factor 1.22) (see Supplementary File 1: Duplication factor). Instead, we used the number of unique molecules in each library as determined by quantitative PCR and the proportion of mapped sequences to estimate the total genomic coverage within each library^44^ as a predictor of library complexity (see Methods: Calculations). The range of estimated genomic coverages within each sampling location was asymmetrically distributed and the data were subsequently transformed by a factor of X^0.1^ in order to fit a linear model, as suggested by Box-Cox transformation, to evaluate significance (Figure 3, for untransformed data and analysis see Supplementary Figure S11 and Supplementary File 1: Est. genomic coverage). Here, the petrous pyramid has the greatest potential to provide higher genomic coverage from an individual library (untransformed median estimated genomic coverage 501.55x, p-values < 0.0056), where all other sampling locations aside from the cementum were statistically indistinguishable (untransformed median estimated genomic coverages for each sampling location: 74.54x for the vertebral body, 55.94x for the phalanx, 46.51x for the pulp chamber, 41.44x for the talus, 17.38x for the superior vertebral arch, and 7.14x for dentin). DNA libraries derived from cementum yielded significantly lower estimates of genomic coverage within each library compared to all other sampling locations (untransformed median of 10.42x, p-values < 0.047) except for those libraries from dentin and the superior vertebral arch (Figure 3). Normalized for input material, cementum yielded similarly low average genomic coverage (0.63x per mg input) while material from the dental pulp chambers yielded the highest (14.98x per mg input), followed by the petrous pyramid (9.44x per mg input) (see Supplementary Figure S15, Supplementary File 1: Est. genomic coverage/mg). The ratio of nuclear to mitochondrial reads (as calculated from mapping to the hg19 genome) had a similarly asymmetrical distribution within sampling locations and as such was transformed by a factor of X^0.5^ to fit our model (for untransformed data see Supplementary Figure S12). We find that nuclear reads were lowest in dentin (untransformed median 1:2769, p-values < 0.011), followed by the pulp chamber (untransformed median 1:539 and not significant when compared to cementum, p-value > 0.45), with all other sampling locations statistically indistinguishable (individual untransformed medians 1:64 in the vertebral body, 1:94 for the distal phalanx, 1:109.86 in the petrous pyramid, 1:128 in the superior vertebral arch, and 1:246 in the cementum) (Figure 4, Supplementary File 1: MT/Nuclear).

**Figure 3.**
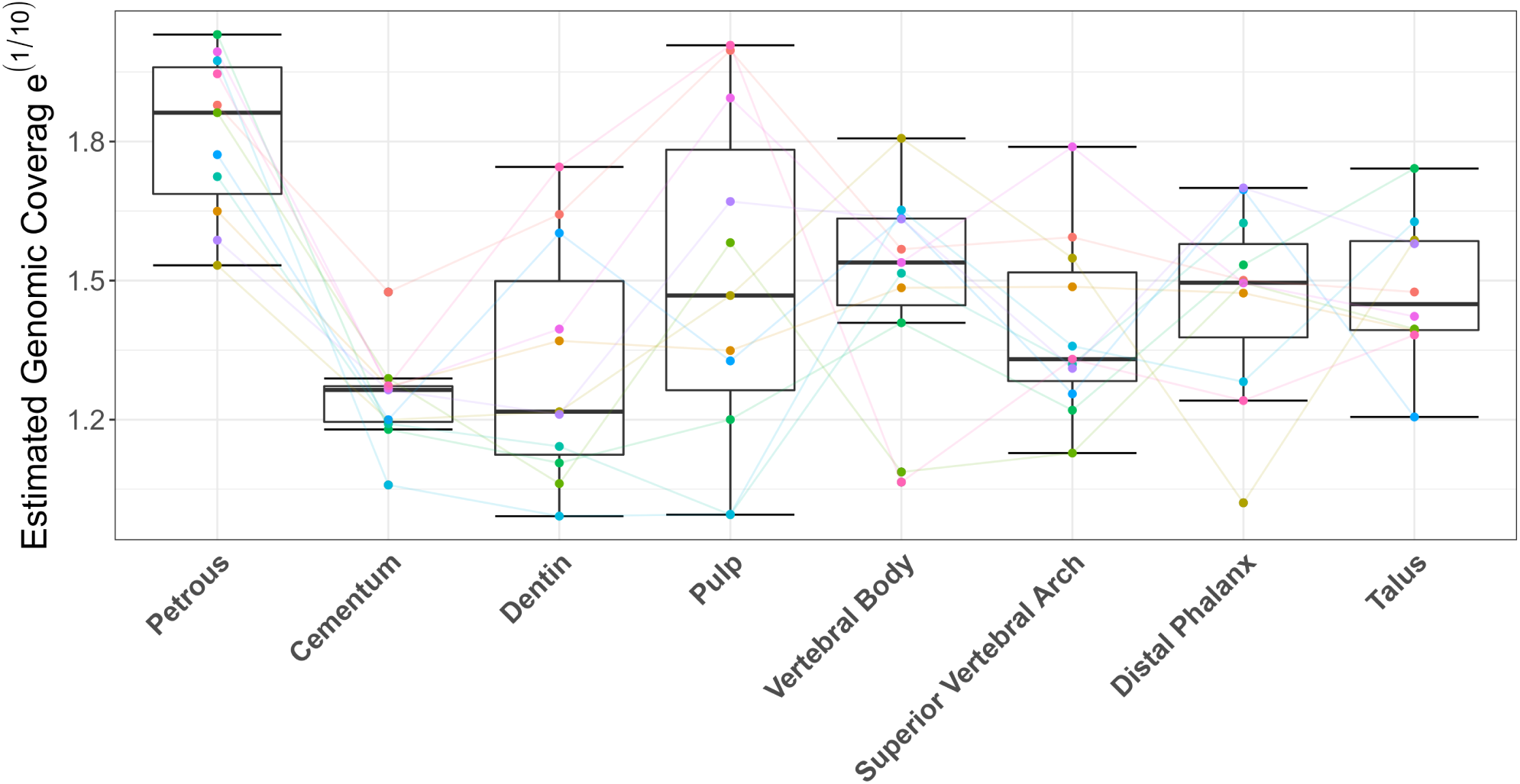
Estimated coverage of the hg19 reference genome contained within each single-stranded library (transformed to the power of X^0.1^ as suggested by Box-Cox transformation for the purposes of fitting the mixed effect model). Coloured points and lines denote sampling across individuals.

**Figure 4.**
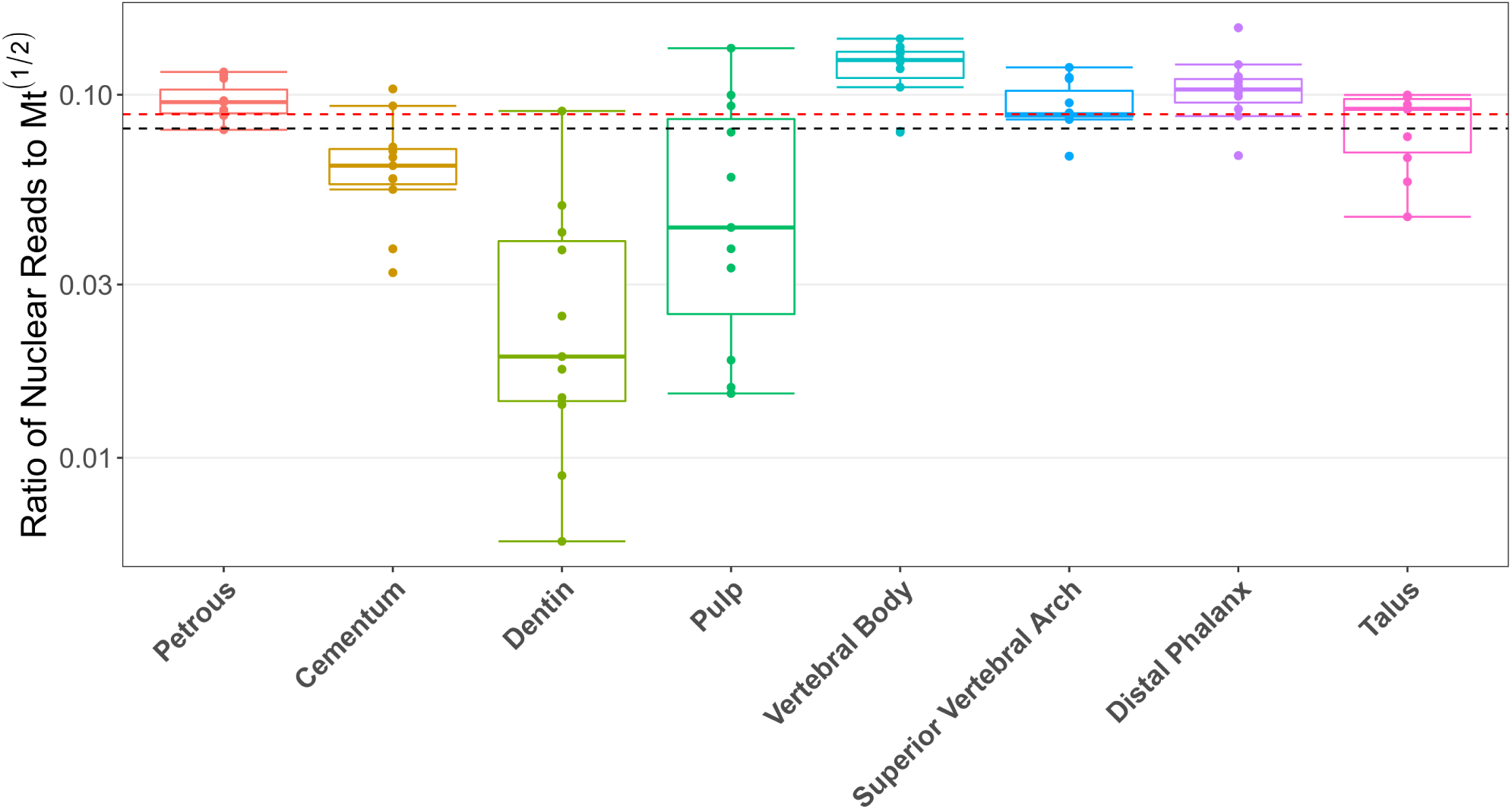
Ratio of reads originating from the mitochondria to those of the nuclear genome (transformed by X^-.05^ as suggested by Box-Cox transformation for the purpose of fitting the mixed effect model). The black line denotes the overall average, the red the overall median.

Contamination estimates based on X chromosome mapping coverage were calculated for all enriched libraries originating from individuals genetically assigned as male (n=7, 8 samples per individual, 56 total samples) using the ANGSD pipeline^45^ to scan known informative SNPs on the X chromosome for polymorphisms. All but one of the 56 samples exhibited low contamination with values statistically indistinguishable across sampling locations (< 4% X chromosome contamination for all enriched libraries from all sampling locations other than the superior vertebral arch of individual five (KRA005), which exhibited contamination levels of 19.52%; p-value = 0.48; see Table 2). Regarding contamination estimates, while those derived from mitochondrial data only are useful for estimating contamination in genetically female individuals, they also offer similar accuracy to X chromosome contamination estimates for samples where the nuclear to mitochondrial read ratio exceeds 1:200^46^. As the samples in this study were not sequenced deeply enough to provide accurate mitochondrial contamination estimates, nor were they target-enriched for mitochondrial reads, mitochondrial contamination estimates were not generated for this dataset.

**Table 2.** Duplication levels, average fragment length, and X chromosome contamination estimates for top performing sampling locations.

| **Sampling Location** | **Average Cluster Factor (#All Mapping Reads/#Unique Reads) Pre-Enrichment (Post-Enrichment)** | **Average Fragment Length (Median) in bp** | **Contamination Estimates (X chromosome; Average Proportion of Human DNA)** | **Average Number of SNPs Covered on X at $\geq 3x$ (per Million Reads)** |
| --- | --- | --- | --- | --- |
| **Petrous pyramid** | 1.188 (1.159) | 65.40 (60.09) | 0 | 73.83 |
| **Cementum** | 1.197 (1.288) | 67.28 (61.36) | 0.011 | 94.78 |
| **Dentin** | 1.188 (1.283) | 60.22 (55.54) | 0.002 | 57.33 |
| **Pulp** | 1.179 (1.206) | 55.14 (50.55) | 0.013 | 44.88 |
| **Distal Phalanx** | 1.191 (1.257) | 65.95 (59.36) | 0.013 | 127.75 |
| **Vertebral Body** | 1.194 (1.247) | 66.14 (60.54) | 0.008 | 119.71 |
| **Superior Vertebral Arch** | 1.190 (1.208) | 63.02 (57.91) | 0.021* | 51.13 |
| **Talus** | 1.198 (1.206) | 68.20 (62.40) | 0.011 | 92.50 |

\*the sample from KRA005 removed as an outlier with a very high (.195) contamination estimate.

Read length and deamination patterns were also evaluated across the eight sampling locations with the highest average human DNA proportions. After filtering to remove all reads < 30bp, the dental pulp chamber housed significantly shorter reads in comparison to all other sampling locations except for dentin (averages of ca. 55bp and 60bp respectively in comparison to the overall average of 63.92bp, p-values < 0.019) (Table 2, Supplementary File 1: Average length). Additionally, we find significant variation in both the frequency of C→T damage caused by nucleotide misincorporations at the ends of the reads and how far into the reads this signal can be detected (Figure 5, Supplementary File 1: Damage signals). Within sampling locations, variations in the frequency of C→T damage patterns were very low (Supplementary Figure: S13, Supplementary File 1: Damage signals), suggesting that the variations observed across sampling locations are unlikely to be due to contamination from modern human DNA. Reads generated from the petrous pyramid have the highest damage signal, a 5’ terminal C→T frequency of ca. 21% on average (all pairwise comparison p-values < 0.001). By comparison, cementum shows significantly lower signals than all other sampling locations (all pairwise comparison p-values <0.001), with approximately half this frequency of damage at the terminal 5’ position. The distal phalanx, talus, and vertebral body form a statistically indistinguishable group with deamination frequencies slightly higher on average compared to the cementum, followed by the dentin, the dental pulp chamber, and the superior vertebral notch, with deamination frequencies lower than the petrous pyramid but higher than the aforementioned group (all pairwise comparison p-values between groupings < 0.001). Average GC content was calculated for all libraries from the eight sampling locations with average human DNA proportions higher than the mean (8.16%) and ranged between 37.14% and 39.87% (see Supplementary File 1; GC content).

**Figure 5.**
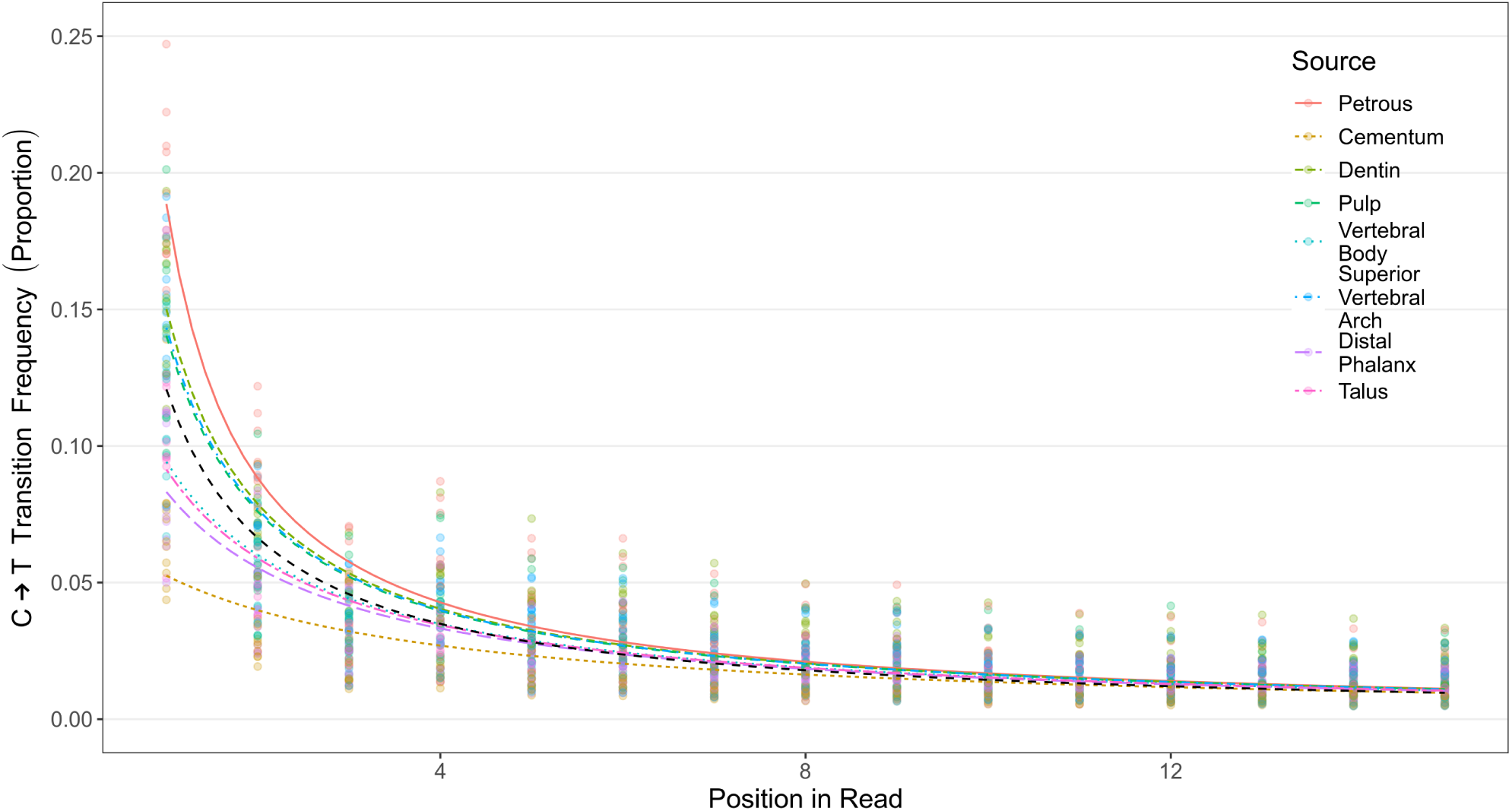
Average proportion of C→T transitions as observed in the first 15 reads of the 5’ end of our reads. The black line represents the mean damage observed across all elements and individuals. Coloured lines indicate the average proportion of transitions within sampling locations, while points represent the corresponding range of individual data within each sampling location.

Since many aDNA analyses, especially those used in population genetics, require a relatively high coverage of informative loci across the genome, libraries are often enriched for these loci by targeted-capture. In our case, this was done for the eight sampling locations that yielded human DNA in proportions higher than the calculated mean for our dataset. To determine the practical usability of the data generated, we compared the relative number of SNPs covered by at least two reads (per million reads sequencing depth) post-1240k capture-enrichment across these eight sampling locations. Here we find that SNP coverage per million reads sequencing effort is statistically indistinguishable between sampling locations. Given that these libraries were not sequenced to exhaustion, this strongly suggests all of these sampling locations are equally suited for SNP analyses at our current sequencing depths (Figure 6). When normalized for available input material the cementum provided significantly higher SNP coverage than all other sampling locations (p-values < 0.02) (See Supplementary Figure S16). As an alternative example of practical usability, we also investigated the phylogenetic resolution for Y-haplotype assignment among all seven male individuals using the ISOGG list of diagnostic SNPs (current as of 26 November 2019) to determine how confidently Y-haplogroups could be called at the ca. 40 million read sequencing depth considered here. The resolution of Y-haplotype assignment was high across most elements and individuals (Table 3). In two individuals (KRA003 and KRA004), the dentin and pulp chamber had a much lower resolution compared to other elements; however, this is most likely an artefact of the low human DNA proportions observed in these samples both before and after SNP capture (Supplementary File 1: % mapping q37, Sheets 1 and 2 respectively), rather than any biological trend.

**Table 3.**
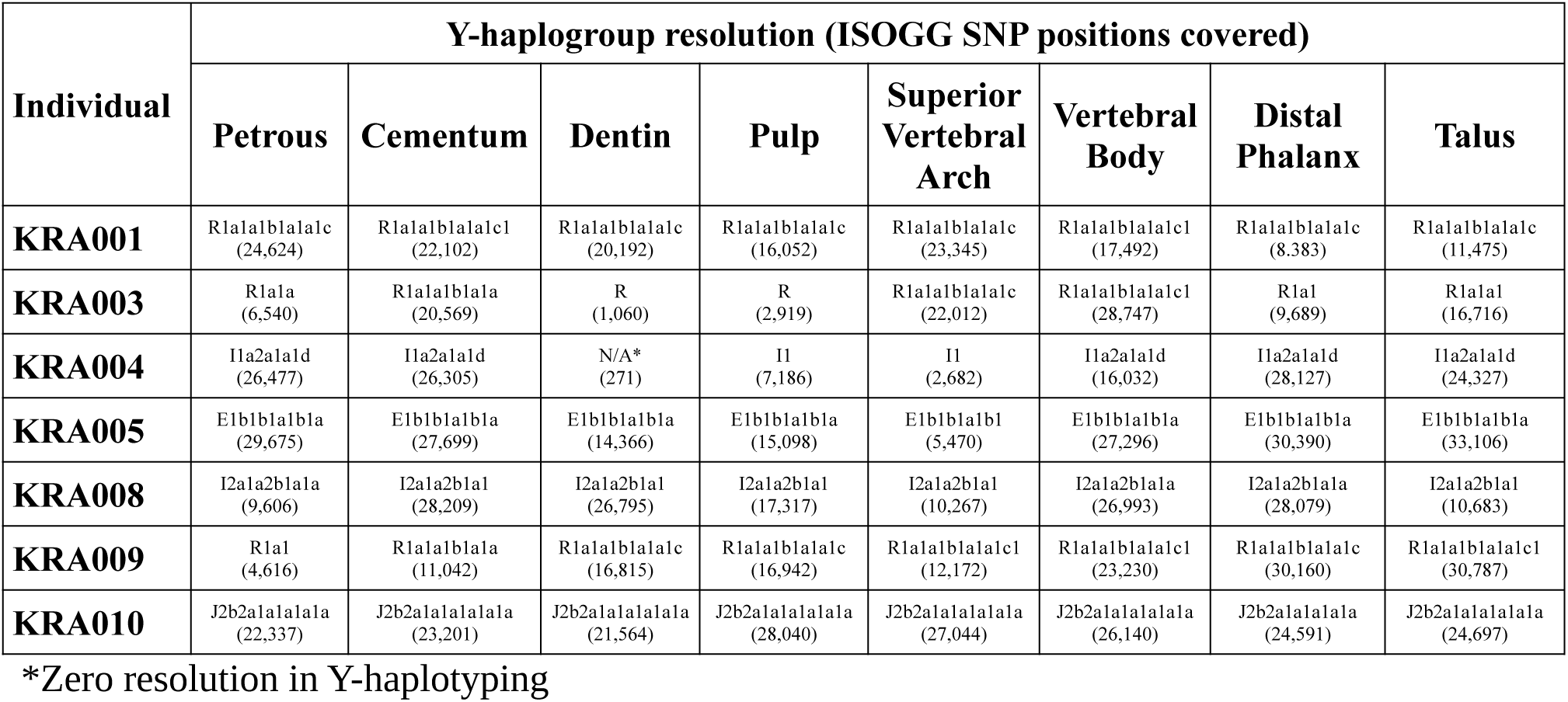
Y-haplotyping resolution post-1240k enrichment across all males and associated sampling locations.

**Figure 6.**
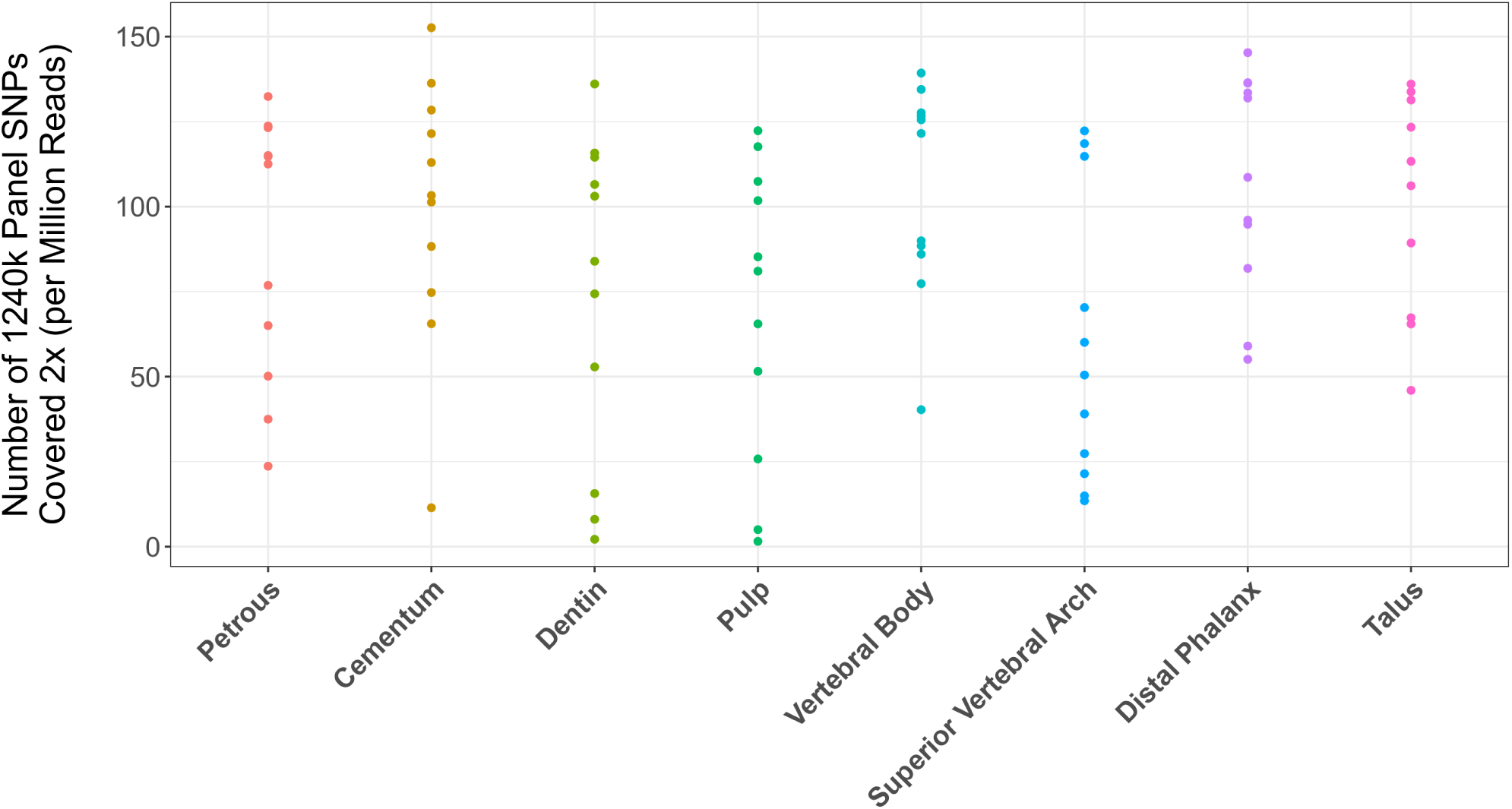
Comparison of 1240k SNP positions covered at least 2x post-capture across skeletal elements normalized by sequencing effort (number of raw reads generated) shown in SNPs per million reads generated.

## Discussion

The extraction of ancient endogenous DNA from archaeological human remains is a labour-intensive and costly process that often yields relatively small amounts of useful data compared to modern specimens^33^. Despite this, when protocols and analyses are optimized, these efforts can nonetheless result in important discoveries about human history, such as determining the genetic relationships between extinct hominins based on whole genome data^4,47,48^, and investigating the correlation between the dissemination of cultural innovations and human migrations^49,50^. The exploration of this type of genetic information can only be achieved via physical damage to precious, irreplaceable, archaeological specimens. As the demand for large sample sizes in aDNA datasets^51–53^ continues to grow, there is imminent need to investigate approaches that maximise the amount of retrievable genetic information via optimisation of methods while imparting the least possible damage to specimens.

Based on previous successes in DNA recovery, the petrous pyramid is currently the most sought-after skeletal element for aDNA analyses^21–25,40,41^. Our investigation of multiple skeletal elements further confirms the value of the petrous pyramid in the recovery of ancient human DNA (Figure 2A-C). We also find that aDNA libraries constructed from material retrieved from the cochlear region of the petrous pyramid are higher in complexity (in terms of the estimated genomic coverage within each library) than those stemming from all other tested sampling locations (Figure 3) in line with previous studies^26,40,54^. Importantly, however, libraries stemming from the petrous pyramid performed comparably to those from all other sampling locations in terms of fragment length, nuclear reads returned (Table 2, Figure 4, Supplementary File 1: Avg. length and MT reads/Nuclear reads), X chromosome contamination estimates (the lowest of all sampling locations with an average of 0, though not statistically significant, Table 2), and SNP coverage post-1240k enrichment (Figure 6). Human DNA fragments recovered from the petrous pyramid show a much higher frequency of cytosine deamination than any other element^40^ (Figure 5, Supplementary Figure S13, Supplementary File 1: Damage signals), which helps to support their authenticity as ancient^39,55–59^. This high damage may necessitate the removal of damaged bases by read trimming to improve mapping quality, especially in those datasets produced from libraries not treated with repair enzymes such as uracil-DNA glycosylase^60^. While the comparatively lower deamination signal identified in the other sampling locations here may result from modern DNA contamination, our data shows no overall correlation between the proportion of human DNA recovered and the proportion of terminal cytosine deamination. Additionally, we do not observe higher amounts of contamination in other sampling sites based on our X chromosome contamination analysis (Table 2), nor do we see significant variation in deamination patterns within sampling locations across individuals (Supplementary Figure S14). However, a high overall fragment length in conjunction with low deamination frequencies (as observed in cementum) may be indicative of contamination with modern human DNA^61^. A previous comparison of deamination patterns in cementum and petrous pyramid yielded a similar differential to what we report here^40^, where cementum exhibited approximately half the frequency of deamination at the 5’ terminus with no indication of modern contamination. Despite its excellent potential for human aDNA recovery, sampling from the petrous pyramid may not always be possible for a variety of reasons including hesitancy on the part of curators in regards to potential damage to the anthropological record, despite the fact that in cases where skulls are fully preserved and sampling of the temporal bone would otherwise be particularly damaging, cranial base drilling techniques have recently been investigated and recommended^26^.

In the remaining skeletal elements where higher than average proportions of human DNA were recovered (> 8.16%), we find that *in situ* molars are inferred to have a high probability of endogenous DNA recovery across all three separate sampling locations (Figure 2A-C). Library complexity was high in both the dentin and material from the pulp chamber (Figure 3), and contamination estimates low (Table 3). Cementum stands out as having both the highest average fragment length (Table 3) and the lowest deamination frequency (Figure 5), which although consistent with previously mentioned findings^40^ may indicate elevated levels of contamination with modern human DNA, despite a low contamination signals observed in X chromosome analyses (Table 2). The dentin and pulp chamber, conversely, returned the shortest average read lengths and were second only to the petrous pyramid in terms of having the highest proportion of detectable deamination damage. However, the fact that dental samples harbour three sampling locations that performed well in terms of human DNA content and two in terms of post-1240k-capture-coverage is an indication of their value. Our observation that dentin exhibited the lowest post-enrichment coverage out of the top sampling locations could be due to its lower nuclear read to mitochondrial read ratio and thus has fewer nuclear reads in the library available for capture. Of note, despite drilling from multiple locations, the enamel, which is often examined in isotope^62,63^, histological^64^ and morphological^65,66^ studies, remained entirely unharmed throughout the sampling process. Additionally, minimally invasive sampling methods for teeth have long been established^67^, with avoidance of alterations to enamel structures. Finally, the two sampling locations most limited in available material (in the context of sampling efforts from a single element) are the cementum and the dental pulp chamber. Both of these sampling locations performed well when directly compared to all other sampling locations (with up to 10x more material available for DNA extraction in some cases, Supplementary File 1) regardless of the amount of material used in extraction. When weight of the sample used for extraction is factored in, however, material from the dental pulp chamber and cementum outperforms all sampling locations other than the petrous pyramid with respect to average number of unique reads mapped per mg of input material (Supplementary Section 2.4). This suggests both sampling materials are particularly rich in DNA content though the complexity of this content in the cementum may not be as high as that found in material from the dental pulp chamber. These factors, combined with the known potential for teeth to harbour oral bacterial and pathogen DNA^34,67–70^, make sampling from molars valuable as an alternative to the petrous pyramid.

Two sampling locations on the thoracic vertebrae, namely the cortical bone collected from the vertebral body and the junction of the lamellae and spinous process (the superior neural arch) were found to yield high average proportions of human DNA (Figure 2A-C, Supplementary File 1: % mapping q37 and Unique reads/million). Additionally, library complexity (Figure 3, Supplementary File 1: Est. genomic coverage), average fragment length (Table 2, Supplementary File 1: Avg. length), post-capture SNP coverage (Figure 6), nuclear to mitochondrial read ratio (Figure 4, Supplementary File 1: MT/Nuclear), and deamination frequencies (Figure 5, Supplementary File 1: Damage signals) fell well within the ranges of the other top performing sampling locations (aside from the petrous pyramid). As with teeth, thoracic vertebrae have multiple high-yield sampling sites, are often well preserved, have been shown to harbour traces of ancient pathogens such as tuberoculosis^71,72^, and in the absence of pathological changes, are of less value in morphological studies given that they are numerous.

Both the talus and distal phalanx exhibited high human DNA recovery rates (Figure 2A-C, Supplementary File 1: % mapping q37 and Unique reads/million) and showed high average fragment length (Table 2, Supplementary File 1: Avg. length) and complexity (Figure 3, Supplementary File 1: Est. genomic coverage), as well as low contamination estimates (Table 2), nuclear-mitochondrial read ratios (Figure 4, Supplementary File 1: MT/Nuclear), and deamination frequency at the 5’ terminus (Figure 5, Supplementary File 1: Damage signals). While both elements have been under-utilised in aDNA investigations to date, the distal phalanx has previously been shown to yield sufficient aDNA to reconstruct a 30-fold genome from a Denisovan specimen^4^.

Among the other sampling locations considered in this survey, those yielding human DNA proportions that are, on average, lower than the overall mean (8.16%) were not considered for further analyses, as our goal was to ascertain the most efficient and cost-effective sampling locations from which to retrieve human DNA. As such, we determined that samples from the femur, metacarpal, ischial tuberosity, metacarpal, ribs, and *clavicula*, as well as any samples derived from cancellous (spongy) material (in order of decreasing yield) are all unlikely to yield high amounts of endogenous human DNA. In light of this, we feel sampling from these elements for DNA analysis should be avoided if possible to circumvent the needless destruction of archaeological samples for minimal gains.

## Conclusions

As intensifying ethical scrutiny surrounds the field of aDNA with regards to the destruction of irreplaceable archaeological human remains^27,37,73–75^, it is imperative for those conducting such research to maximize the chances of successful data generation from minimally invasive sampling. It is of similar importance to both maximize the potential amount of information obtained from and to simultaneously minimize laboratory processing times for each sampling effort to balance the high cost of aDNA research with the aforementioned ethical considerations. As such, our large cross-sectional evaluation of aDNA recovery cross the skeleton helps to facilitate this balance by increasing perspectives on molecular preservation not only in previously studied sampling locations, but also in a set of new ones. Our results demonstrate that, from the locations we consider here, the dense inner portion of the petrous pyramid remains the best sampling location for high-quality ancient DNA. However, we also report on seven alternative sampling locations on four other skeletal elements, all of which performed equally well in our evaluation, that show promise as suitable alternatives to the petrous pyramid. Though our sample set is limited both temporally and geographically, our results are likely informative for other climatic regions, time periods and perhaps even in anatomically comparable species as has already been demonstrated for the petrous portions itself^76–79^. It should also be noted that, as this study has focused on identifying the most efficient sampling locations from which host (in this case human) DNA can be recovered, the sampling strategies and suggestions put forth here may not be applicable in studies seeking to retrieve DNA from pathogens, the microbiome, or other co-cohabitating organisms within the host.

By providing researchers with more varied options for the successful recovery of endogenous ancient human DNA, we hope to provide a framework in which successful collaborations between archaeologists and geneticists can continue to enrich our knowledge of history and heritage. At the same time, continuing efforts to fully optimize our sampling strategies will allow the above collaborations to go forward in a more ethical fashion by minimizing damage to the finite archaeological record.

## Methods

### Sample selection, pre-treatment, and bone powder generation

Individuals from the Krakauer Berg collection housed at the State Office for Heritage Management and Archaeology, Saxony-Anhalt (State Museum of Prehistory, Halle (Saale)) (Figure 1) were sampled for DNA extraction. This collection consists of approximately 800 individuals and represents a typical medieval burial, with age and sex distribution consistent with an attritional context. Ten skeletal elements were selected as targets for aDNA sampling (Table 1, Supplementary Material: Section 1.2). For each individual, morphological preservation of these pre-selected elements was assessed, and individuals were included in the study if a minimum of eight elements were present and were sufficiently well preserved. This resulted in a study set of eleven individuals, seven males and four females (genetically assigned, see below), who ranged in age at death from ca. 10-45 years, with two juveniles and nine adults. Radiocarbon dating of ribs from each individual (performed at the Curt Engelhorn Centre for Archaeometry in Mannheim, Germany) placed the skeletal series in a time interval of ca. 1050-1402 cal AD (Table 4).

**Table 4.** Biological sex (genetically determined), age at death (archaeologically determined), and calibrated ^14^C dates (years BP) of individuals selected for aDNA sampling.

| Individual<br>(Laboratory ID) | Archaeological ID<br>(Burial Nr.-Individual Nr.) | Sex | Age At Death | <sup>14</sup>C dates<br>(AD, Cal 2-sigma) |
| --- | --- | --- | --- | --- |
| KRA001 | 25-1a | Male | 25-35 | 1058-1219 |
| KRA002 | 20-2a | Female | 20-22 | 1227-1283 |
| KRA003 | 113-6a | Male | 25 | 1059-1223 |
| KRA004 | 246-1a | Male | 15 | 1284-1392 |
| KRA005 | 276-2a | Male | 10-12 | 1170-1258 |
| KRA006 | 307-4a | Female | 30-40 | 1218-1266 |
| KRA007 | 377-6a | Female | 25-30 | 1167-1251 |
| KRA008 | 436-6a | Male | 20 | 1301-1402 |
| KRA009 | 566-3a | Male | Unknown Adult | 1158-1254 |
| KRA010 | 600-7a | Male | 25 | 1276-1383 |
| KRA011 | 632-2a | Female | 30-45 | 1040-1159 |

To reduce external contamination as much as possible, all elements were processed in a dedicated ancient DNA laboratory under controlled conditions. At least two sampling locations (Table 1, Supplementary Material: Section 1.2) were selected for each element other than the petrous pyramid, one of which was comprised of cortical bone and the other of cancellous bone. Sampling of the petrous pyramid followed previously established sampling procedures^43^ and involved the sectioning of the petrous pyramid to allow access to the dense bone surrounding the cochlea for drilling. Sampling of teeth was performed in a three-step process and involved removal of the cementum followed by sectioning and drilling of the pulp chamber and dentin portions. Prior to sampling, all relevant locations on each element were cleaned with bleach (0.01% v/v) via 5-minute incubation, followed by rinsing with distilled water and exposure to UV light for 30 minutes to cross-link any residual surface contamination from modern DNA. Where applicable the outermost surface of bone was removed by abrasion with a standard dental drill (KaVo K-POWERgrip EWL 4941) and size 016 round bit (NTI Kahla). Approximately 100mg of bone powder was drilled from each sampling location with exception of the cementum and dental pulp chambers where and average of ca. 19mg (standard deviation of 10.8mg) and ca. 24mg (standard deviation of 15.03mg), respectively, of bone powder was recovered, the entirety of which was used for DNA extraction. An average of ca. 54mg (standard deviation of 11mg) of bone powder was used in downstream DNA extractions for all other sampling locations (Supplementary File 1: mg input). For molars, cementum was removed by abrasion using a diamond coated rotary cutting disc (NTI Kahla). The tooth was then sectioned at the cemento-enamel junction using a jeweller’s saw (Präzisions-Sägebogen Antilope, with 75mm blade). Powder from a first pass drilling of the pulp chamber was collected before further sampling of the underlying dentin (Supplementary Material: Section 1.2)

### DNA extraction, library preparation, and sequencing

All DNA extractions were conducted in the clean room facility of the Department of Archaeogenetic of the Max Planck Institute for the Science of Human History (MPI-SHH) located in Jena, Germany, using a modified filter column protocol^14^ (Supplementary section 1.3.1). Single-stranded DNA libraries^80^ were prepared from all extracts by automation^81^ using the Agilent Bravo™ liquid handling system at the Max Planck Institute for Evolutionary Anthropology in Leipzig, Germany. Subsequent to initial analysis, libraries from all sampling locations found to have average human DNA content of 8.16% or greater were enriched by bait capture^82^ for regions in the human 1240k^25^ reference dataset. Sequencing was done via a 75bp paired-end kit on an Illumina HiSeq 4000 platform to a depth of ca. 5 million reads for initial screening and to ca. 40 million reads following 1240k capture enrichment.

### Evaluation Criteria

One of the most common metrics for the evaluation of molecular preservation in archaeological remains percentage of endogenous (*i.e.* human) aDNA recovered after sequencing. However, a high percentage of endogenous DNA on its own provides limited information on the utility of a given DNA library for downstream analysis. For example, it is important that both the proportion of human DNA relative to that of potential contaminants as well as the quantity (*e.g.* the number of sequences mapping to the reference as well as the as the proportion of the reference actually covered) of human DNA are high for whole genome sequencing, whereas the quantity alone is the most important criterion when using target enrichment approaches^83^. Beyond this, the integrity of the DNA molecules themselves plays an important role in the downstream mapping of sequencing data^84,85^ as well as playing an important role in the authentication of ancient DNA^39,55–59^. For this reason, we integrated additional measures of data quality into our initial evaluation^86^, including the quantity of recovered human DNA, estimated DNA library complexity (in terms of both sequence duplication levels and total estimated genomic coverage), estimates of modern human DNA contamination, the ratio of nuclear to mitochondrial read recovery, average DNA fragment length, and patterns of deamination observed in reads mapping to the human reference genome. All resulting data was normalized to reflect outcomes expected from equal sequencing efforts (raw number of sequences generated prior to merging, duplicate removal, as well as length and quality filtering) across all samples where appropriate. The aim of our study was to develop a predictive model of DNA recovery based on the relative performance of each sampling location in terms of quality and quantity of recovered human DNA. We, therefore, opted not to normalize our analyses against the amount of sampling input material, despite the restricted amounts available in some locations (see Supplementary Section 2.4 for analyses normalized for starting material).

### Contamination estimates

Contamination estimates were calculated using the ANGSD^45^ software package to examine the probability of foreign X chromosome contamination in samples from male individuals using the post-capture enrichment data sets generated for eight sampling locations with human DNA recovery above 8.16%.

### Mapping

Human DNA content and sequence quality were determined by mapping reads to the hg19 human reference genome (accession number: GCF_000001405.13) using the EAGER^87^ pipeline: BWA^88^ settings: -n set at 0.1 and a mapping quality filter of q37. To assess resolution of the above pipeline in detecting ancient human DNA sequences, we created a simulated dataset based on the hg19 human reference for mapping evaluation and to act as a best-case scenario for comparative purposes. We first cut the reference sequence into fragments of average length and size distribution modelled after a representative sample (KRA001.B0102, petrous pyramid single-stranded library; see Supplementary File 1: Average and Median length). We then used the software Gargammel^89^ to artificially add a deamination pattern to the data that simulated an ancient DNA damage signal consistent with the same sample (see Supplementary File 1: Damage signals). The resulting simulated aDNA dataset was then mapped as above.

### Calculations

Percentage of human reads recovered from each sampling effort was calculated as:

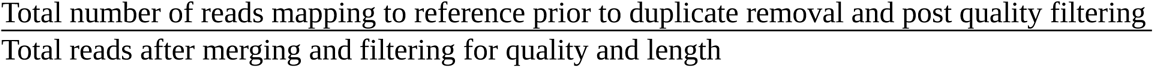

The number of unique reads mapping to the human genome per million reads sequencing effort was calculated as:

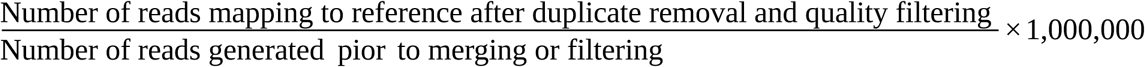

Total genomic coverage within a library^44^ was estimated by calculating:

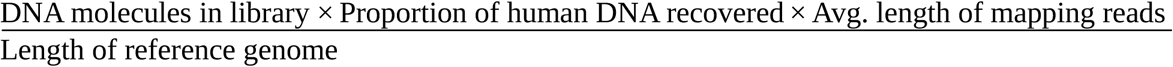

### Mixed Effects Modelling

All statistical analyses involving generalized linear models and mixed effects models described here were performed using the *R Statistical Software Package*^90^, where a p-value of 0.05 was considered significant. When multiple hypotheses were performed, p-values were adjusted to control for a family-wise error rate of 0.05 using the *p.adjust* function.

In all mixed effects models we considered the skeletal element to be a fixed effect with the individual as a random effect. Backward model selection was performed using ANOVA, including for testing whether random effects in the final analyses were deemed significant.

When modelling response variables with an obvious upper bound (*i.e.* endogenous DNA content of 100%), we implemented beta mixed effects regression as implemented in the *glmmTMB* package^91^. Optimal power transformations for theoretically unbounded response variables were performed using a Box-Cox transformation as implemented in the *MASS* package^92^.

We compared the effects of skeletal elements on response variable by inspecting the estimated marginal means in our optimal mixed effects and fixed effects models using the *emmeans* package^93^.

## Acknowledgements

The authors would like to thank the laboratory staff at the Max Planck Institute for Evolutionary Anthropology, Leipzig, Germany as well as all the technicians, students, and scientific colleagues at the Max Planck Institute for the Science of Human History, Jena, Germany, with particular thanks to technicians Antje Wissgott and Franziska Aron for aiding in the laboratory work behind this publication as well as Elizabeth Nelson for her help in identifying osteological features. In addition, the authors would also like to thank the State Office for Heritage Management and Archaeology, Saxony-Anhalt (State Museum of Prehistory, Halle (Saale)) for opening up their collection and providing all samples used in this study and Xandra Dalidowski for leading the excavation. This study was funded by the Max Planck Society, the European Research Council (ERC) under the European Union’s Horizon 2020 research and innovation program under grant agreements No 771234 – PALEoRIDER (WH, ABR) and Starting Grant No. 805268 CoDisEASe (to KIB).

## Author contributions

CP is the primary author and was responsible for the gathering, processing, sampling from, and DNA extraction from all samples, as well as their subsequent analyses. ABR performed all statistical analyses and coding, as well as authoring the corresponding methods sections and the editing of the overall manuscript. SF of the State Office for Heritage Management and Archaeology, Saxony-Anhalt (State Museum of Prehistory, Halle (Saale)) contributed archaeological remains sampled in this study and the archaeological context. SN produced single-stranded libraries for all samples at the Max Planck Institute for Evolutionary Anthropology, Leipzig, Germany. MM oversaw single-stranded library preparation at the Max Planck Institute for Evolutionary Anthropology, Leipzig, Germany, aided in the editing of the manuscript. KB acted as co-supervisor to the primary author, provided funding, aided in the experimental design of this study, and contributed to the writing and editing of this manuscript. WH acted as co-supervisor to the primary author, provided funding, aided in the experimental design of this study, coordinated sample selection, and contributing to the writing and editing manuscript. JK acted as co-supervisor to the primary author, aided in experimental design, and provided funding for the study.

## Code Availability

All programs and R libraries used in this manuscript are freely and publicly available from their respective authors. All custom written R code is available by request.

## Data availability

Sequence data is available through the European Nucleotide Archive under accession number PRJ-EB36983 (released upon publication).

