## Supplementary Material for "A systematic investigation of human DNA preservation in medieval skeletons"

### 1. Laboratory Processing

#### 1.1 Pre-treatment

All samples were initially cleaned with 0.01% v/v bleach to remove dirt, then rinsed with distilled water before being exposed to ultraviolet light for 30 minutes.

#### 1.2 Bone powder generation

All bone powder was generated by drilling using a standard dental drill with standard drill bit on a low-speed, high-torque setting unless otherwise noted.

##### 1.2.1 *Pars Petrosa*

The petrous pyramid was first cut in half along the lateral line using a jeweller's saw (Figure S1A). The interior portion was then visibly examined and bone powder generated from the densest area (Figure S1B).

A.

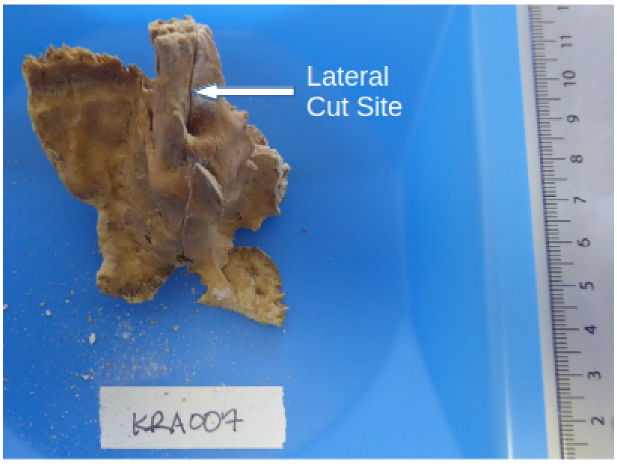


B.

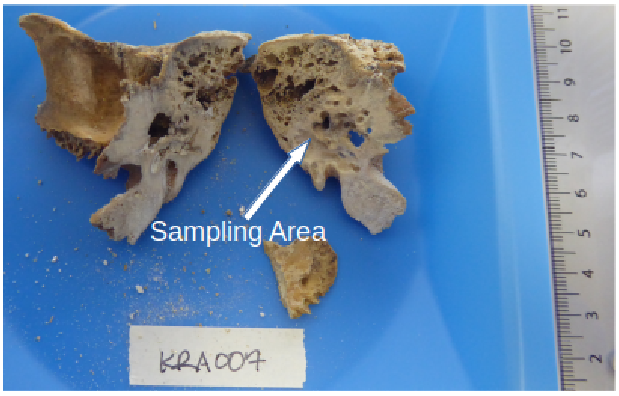


**Figure S1A-B:** *Pars petrosa* pre-sectioning (A) and post showing the sectioning and subsequent drilling sites (B).

#### 1.2.2 Teeth

Cementum was removed from the root portion of the tooth using a standard dental drill fitted with a circular cutting attachment. The blade of the cutting wheel was placed lightly against the root at a 20° angle (relative to the bottom of the root) on a low-speed, high-torque setting and the cementum scraped off downward (Figure S2A-B). The tooth was then bisected along the cementum-enamel junction. Powder from pulp chamber was generated using a standard dental drill bit from the first pass of the interior of the crown. Subsequent passes were used to generate bone powder from dentin (Figure S2C).

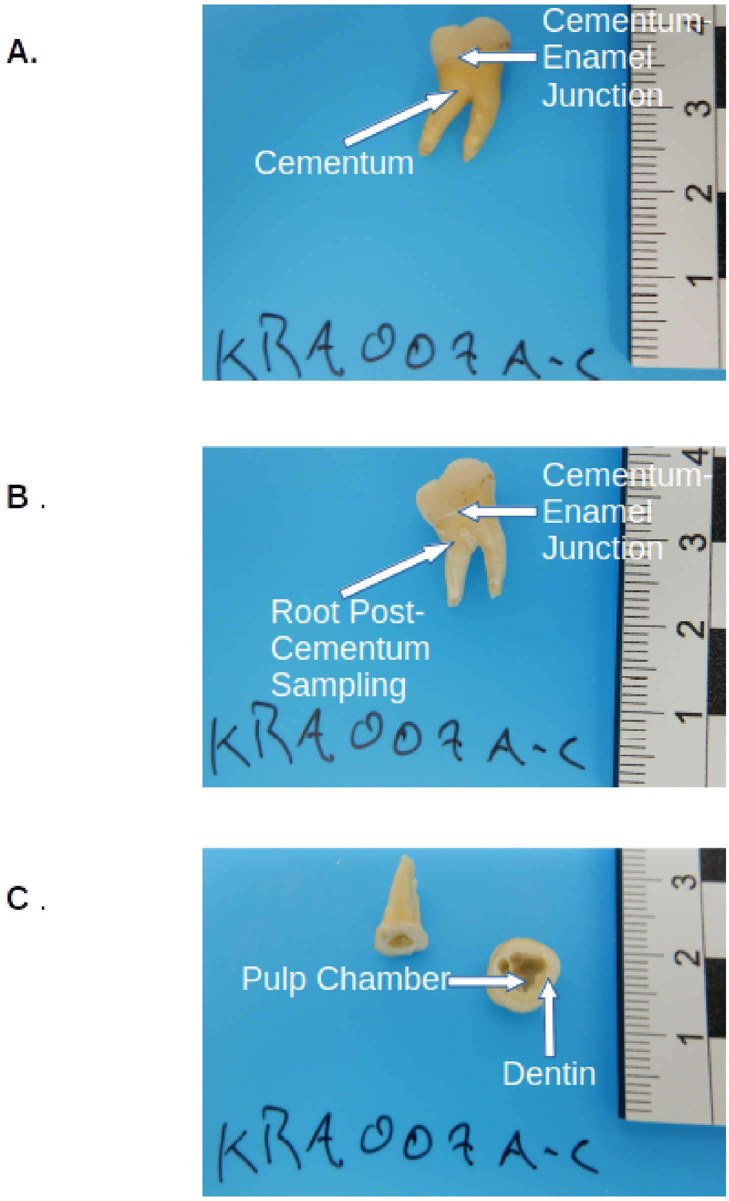


**Figure S2A-C:** *In situ* molar pre (A) and post (B) removal of cementum, as well as pre (B) and post (C) sectioning and drilling of the pulp chamber and underlying dentin.

#### 1.2.3 Clavicles

Cortical bone powder was harvested from the exterior apex of anterior sternal curve of the shaft of the clavicle, and cancellous bone powder from the interior of the acromial facet (Figure 3).

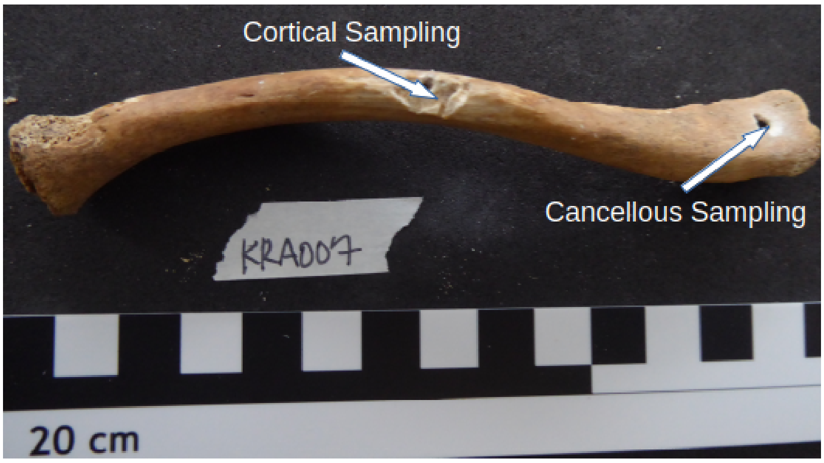


**Figure S3:** Sampling sites of the clavicle showing drilling locations for both cortical and cancellous tissue.

#### 1.2.4 Vertebrae

Cortical bone powder was generated the spine of the spinous process, the exterior of the vertebral body, the interior surface of the neural foramen, and the superior apex of junction of the lamellae and spinous process (superior vertebral arch). Cancellous bone powder was harvested from the interior of the vertebral body (Figure S4).

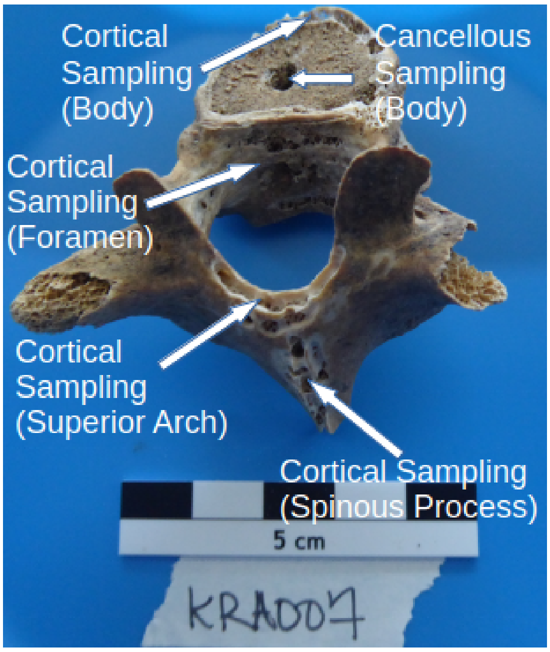


**Figure S4:** All sampling locations (post-drilling) of the thoracic vertebrae.

**1.2.5 Ribs (1st)**

Cortical bone powder was sampled from the outer surface of the serratus anterior, cancellous powder from the interior of the costoclavicular ligament attachment site (Figure S5).

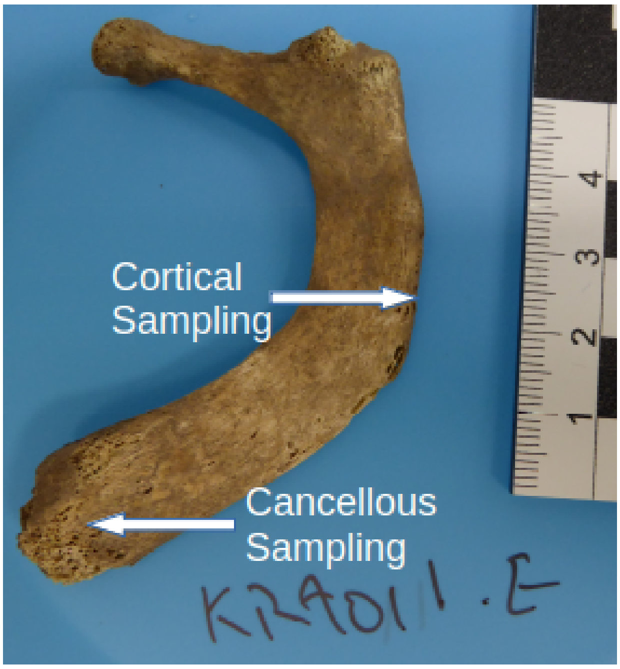


**Figure S5:** Cortical and Cancellous sampling locations on the first rib.

**1.2.6 Metacarpals**

Cortical bone was harvested from the exterior surface of the shaft, cancellous from the interior of the head (Figure S6).

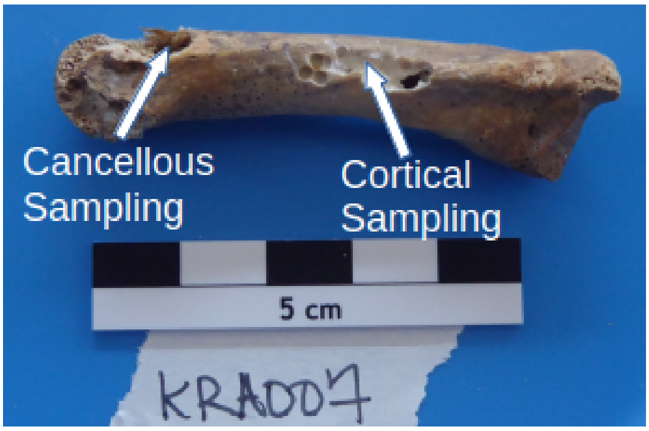


**Figure S6:** Metacarpal showing drilling locations for the collection of both cortical and cancellous tissue

#### 1.2.7 Distal Phalanx

Cortical bone powder was harvested from the pad and shaft of the distal phalanx, cancellous bone from the interior of the base (Figure S7).

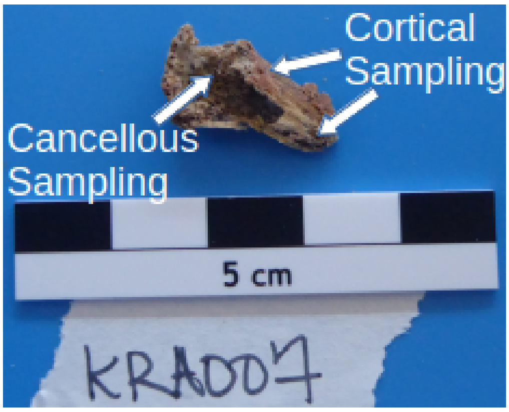


**Figure S7:** Distal phalanx showing drilling locations for the collection of both cortical and cancellous tissue.

#### 1.2.8 Ischial Tuberosity

Cortical bone was harvested from the exterior surface of the tuberosity, cancellous from the interior (Figure S8).

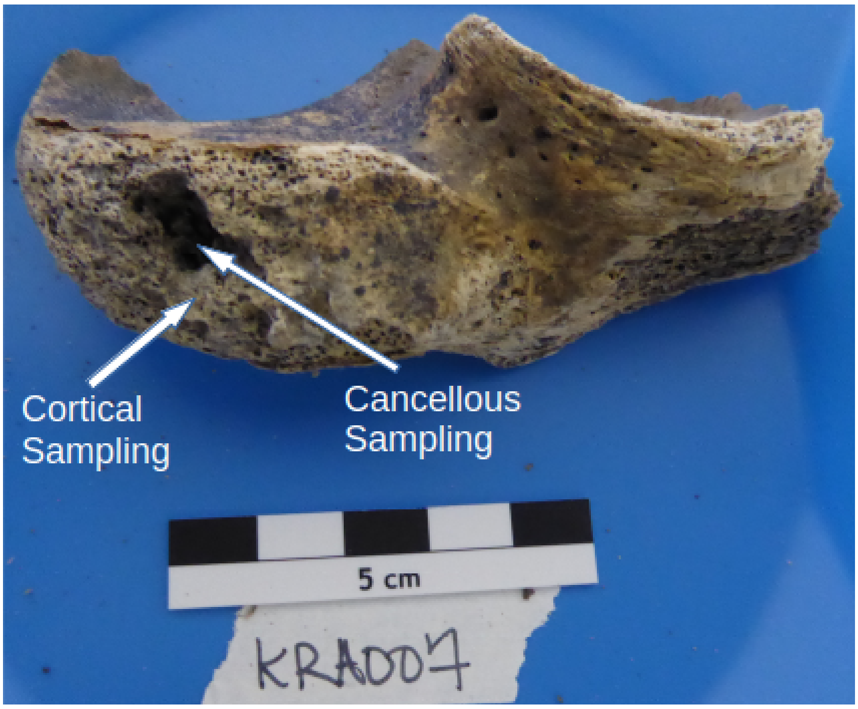


**Figure S8:** Ischial tuberosity showing drilling sites for the collection of both cortical and cancellous material.

#### 1.2.9 Femora

Cortical bone material was harvested from the shaft, just below the lesser trochanter and cancellous from the interior of the head (Figure S9).

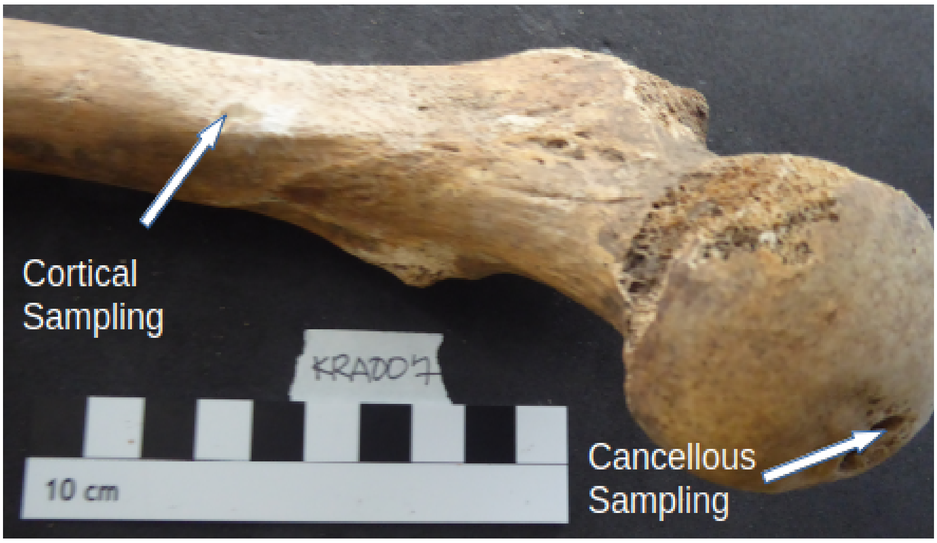


**Figure S9:** Femur showing drilling sites for the collection of both cortical and cancellous material.

#### 1.2.10 Tali

Dense tissue was harvested from the "neck" and articular surface, less compact cancellous from the interior of the medial facet (Figure S10).

A.

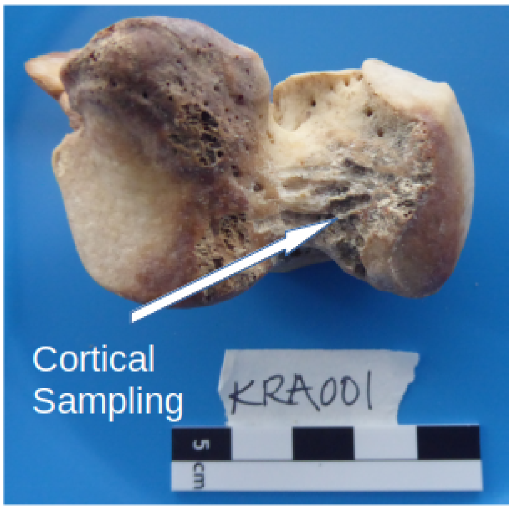


B.

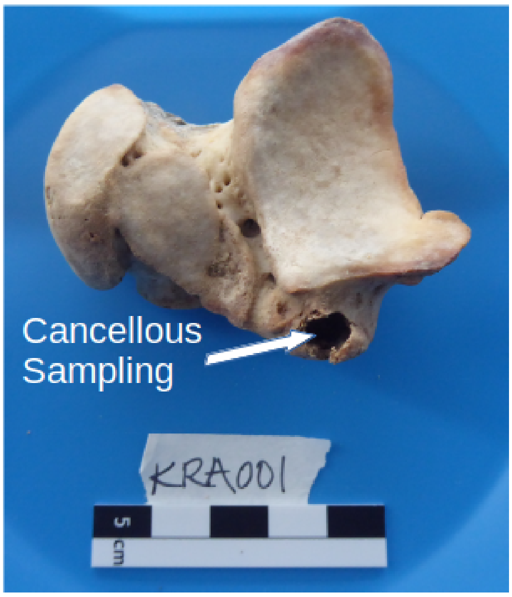


**Figure S10:** Talus showing drilling sites for the collection of both cortical (A) and loosely packed cancellous tissue (B).

#### 1.3 DNA extraction

One millilitre of UV-purified extraction buffer (900µl 0.5M EDTA; 75µl UV-treated, HPLC grade water; and 25µl 10mg/ml Proteinase K) was added to approximately 50mg of bone powder (where available) in a 2ml Eppendorf Biopur microcentrifuge tube and incubated (with rotation) overnight at 37°C. The mixture was then centrifuged for 2 minutes at 14000rpm and the supernatant collected and transferred to 50ml Falcon tube containing 10ml of UV-treated binding buffer (6ml 5M GuHCl, 4ml isopropanol) and 400µl 3M sodium acetate (pH: 5.2) and the contents mixed by inversion. This was then transferred into the funnel of a High Pure Viral Extract Extender Assembly and centrifuged for 8 minutes at 1500rpm. The column was then removed from the funnel and transferred to a fresh collection tube before being dry-centrifuged for 2 minutes at 14000rpm. 450 ml of wash buffer (provided in the High Pure Viral Nucleic Acid Extraction Kit) was then added and the column spun at 14000rpm for 1 minute. The collection tube was then emptied and the wash repeated. The column was then dry-spun at 14000rpm for 1 minute before being transferred to a 1.5ml siliconized microcentrifuge tube for elution. Elution was done in two cycles of adding 50µl of Tris-EDTA-Tween buffer (Applichem low EDTA 1x Tris-EDTA buffer, 0.05% Tween 20), incubating at room temperature for 3 minutes, and centrifuged at 14000rpm for 1 minute, resulting in 100µl of purified aDNA extract.

### 2. Supplemental Analysis

#### 2.1 Expected proportion of human DNA recovered simulations

**Table S1.** Frequency of observed rankings of skeletal elements in terms of human DNA-richness after 55,000 simulated samplings.

| **Skeletal Element** | **Frequency of observed rankings in endogenous DNA-richness (%)** | | | | | | | |
|---|---|---|---|---|---|---|---|---|
| | **1st** | **2nd** | **3rd** | **4th** | **5th** | **6th** | **7th** | **8th** |
| ***Pars petrosa*** | 41.87 | 22.08 | 14.03 | 9.68 | 6.40 | 4.20 | 2.07 | 0.81 |
| **Cementum** | 10.23 | 13.30 | 14.08 | 14.30 | 14.07 | 13.59 | 12.38 | 9.20 |
| **Dentin** | 6.34 | 9.03 | 10.74 | 12.50 | 13.80 | 15.16 | 16.94 | 16.62 |
| **Pulp** | 7.23 | 10.10 | 11.73 | 12.85 | 13.99 | 14.96 | 15.78 | 14.52 |
| **Vertebral Body** | 10.61 | 13.58 | 14.34 | 14.45 | 14.19 | 13.22 | 11.88 | 8.88 |
| **Superior Vertebral Arch** | 4.28 | 6.44 | 8.59 | 10.12 | 12.00 | 14.77 | 19.51 | 25.44 |
| **Distal Phalanx** | 10.65 | 13.91 | 14.43 | 14.30 | 14.02 | 13.48 | 11.92 | 8.42 |
| **Talus** | 9.93 | 12.72 | 13.21 | 12.94 | 12.67 | 11.76 | 10.67 | 8.06 |

**2.2 Untransformed data for both estimated genomic coverage and nuclear to mitochondrial read ratio.**

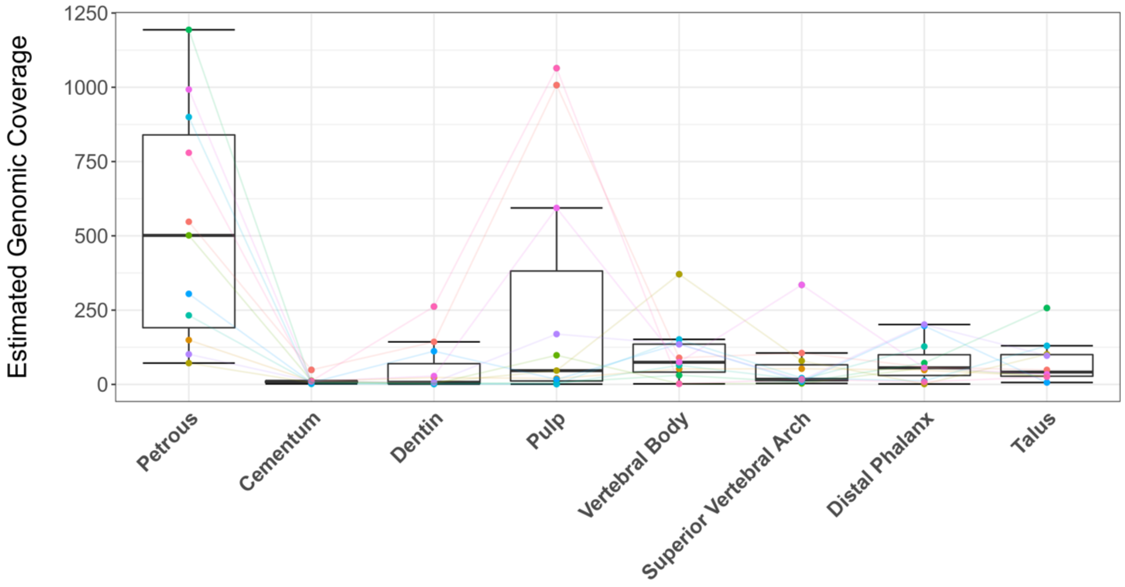


**Figure S11.** Untransformed estimated genomic coverage across the eight sampling locations with average proportion of human DNA content higher than the overall mean (>8.16%). Coloured points and lines represent the genomic coverage across sampling locations within an individual.

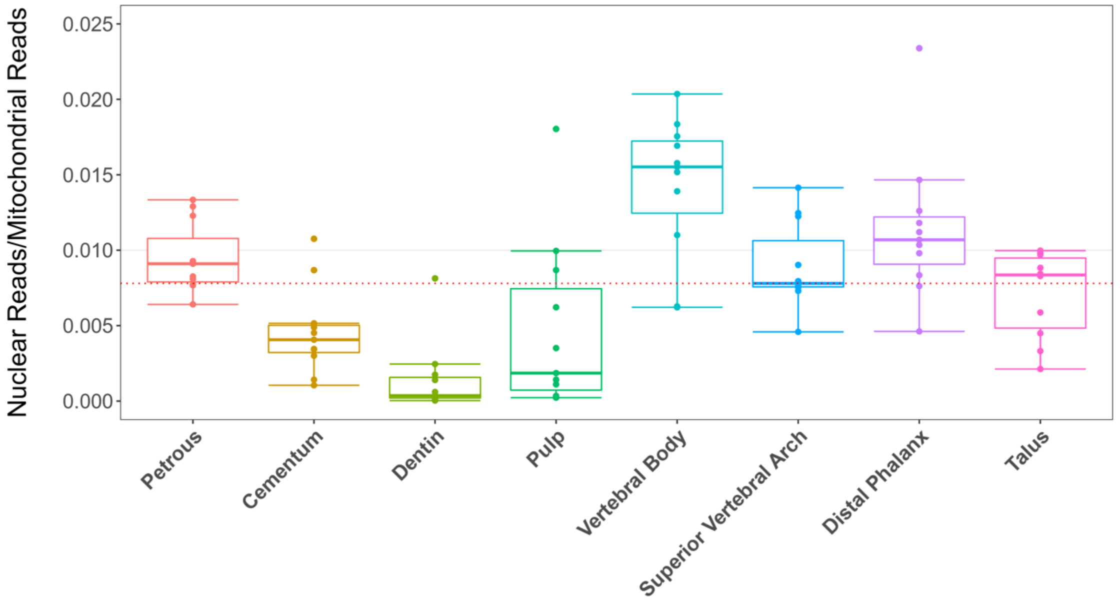


**Figure S12.** Untransformed nuclear to mitochondrial read ratio across the eight sampling locations with average proportion of human DNA content higher than the overall mean (>8.16%). The red line represents the overall median.

### 2.3 Consistency of deamination patterns across both sampling location and individual.

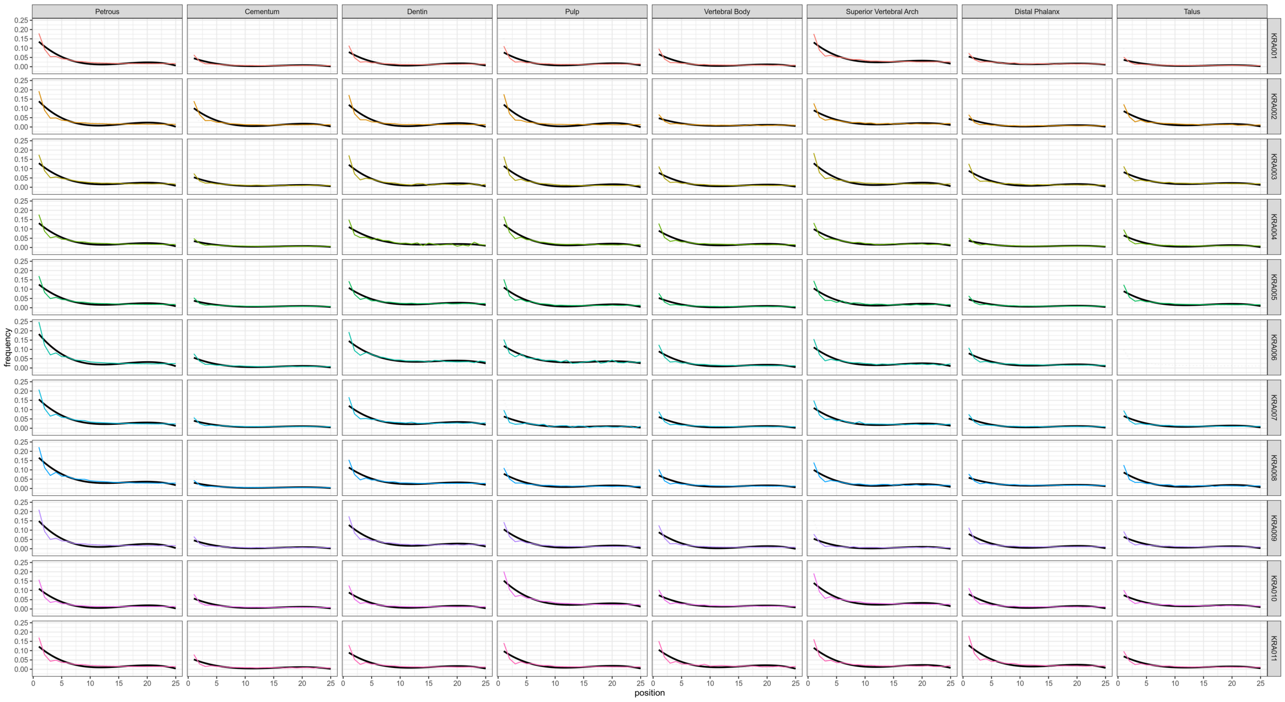


**Figure S13.** Deamination patterns for all eight sampling locations with higher average human DNA content than the overall mean (>8.16%) showing the consistency within each sampling location. Black lines represent lines of best fit.

#### 2.4 Richness per milligram input material

The per mg richness of each individual sampling location is beyond the scope of this investigation. However, it was still noted that the cementum and the dental pulp chambers, despite yielding less starting material, were still comparable to all other sampling locations. As such, all relevant analyses were also performed after normalization of milligrams of material used in each DNA extraction.

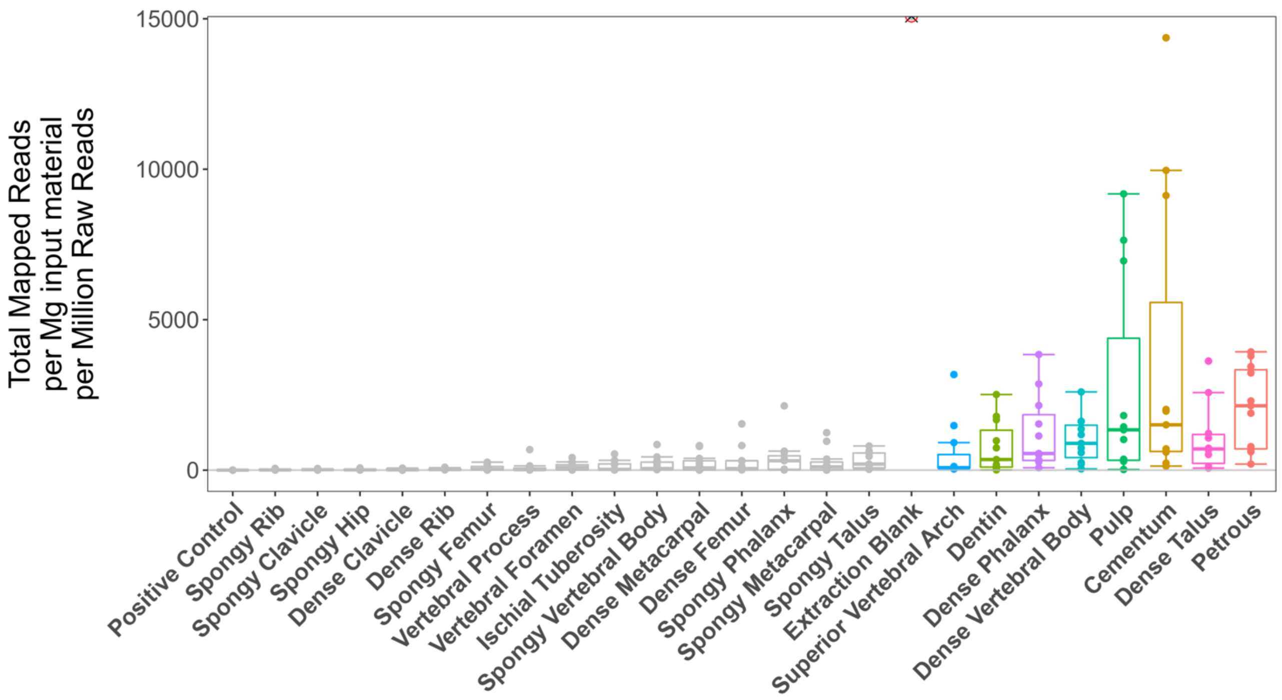


**Figure S14.** Total unique reads mapped to the hg19 human reference after normalization for starting input material and sequencing effort, showing the dental pulp chambers and cementum to be especially rich in DNA per mg.

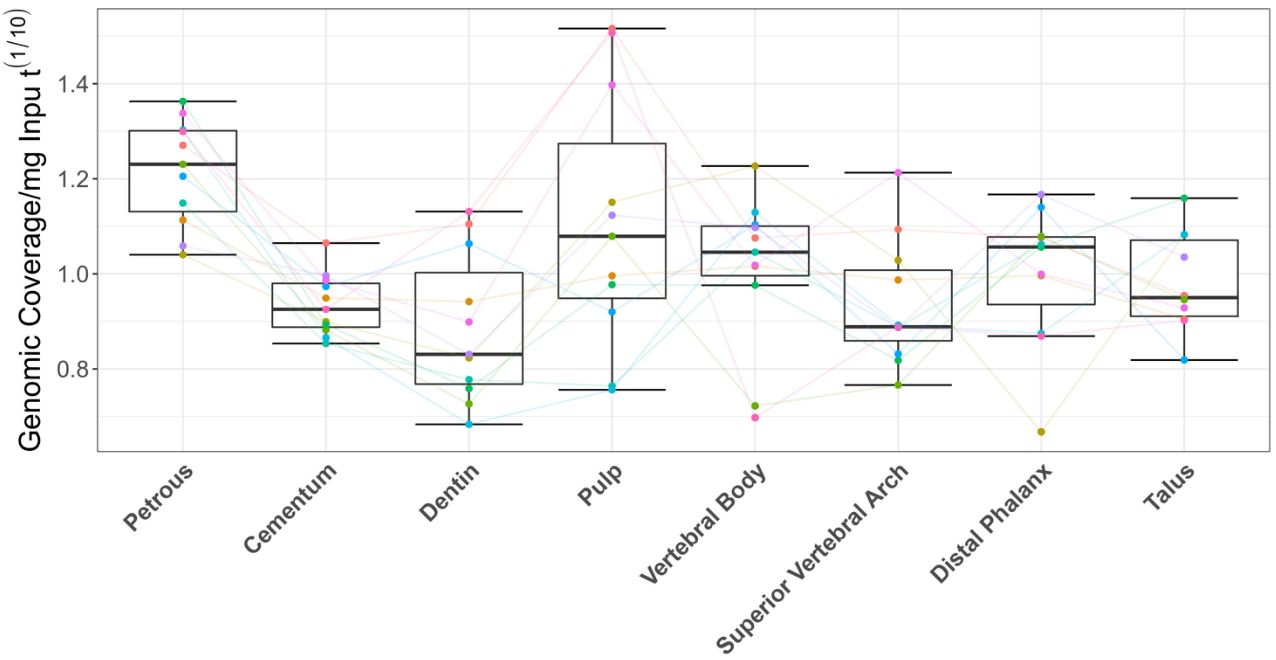


**Figure S15.** Estimated genomic coverage/mg contained within DNA libraries from each sampling location, showing the richness of the dental pulp chamber and increased richness in the cementum (to a level comparable with all other sampling locations) when input material is factored in. Coloured points and lines represent values within individuals.

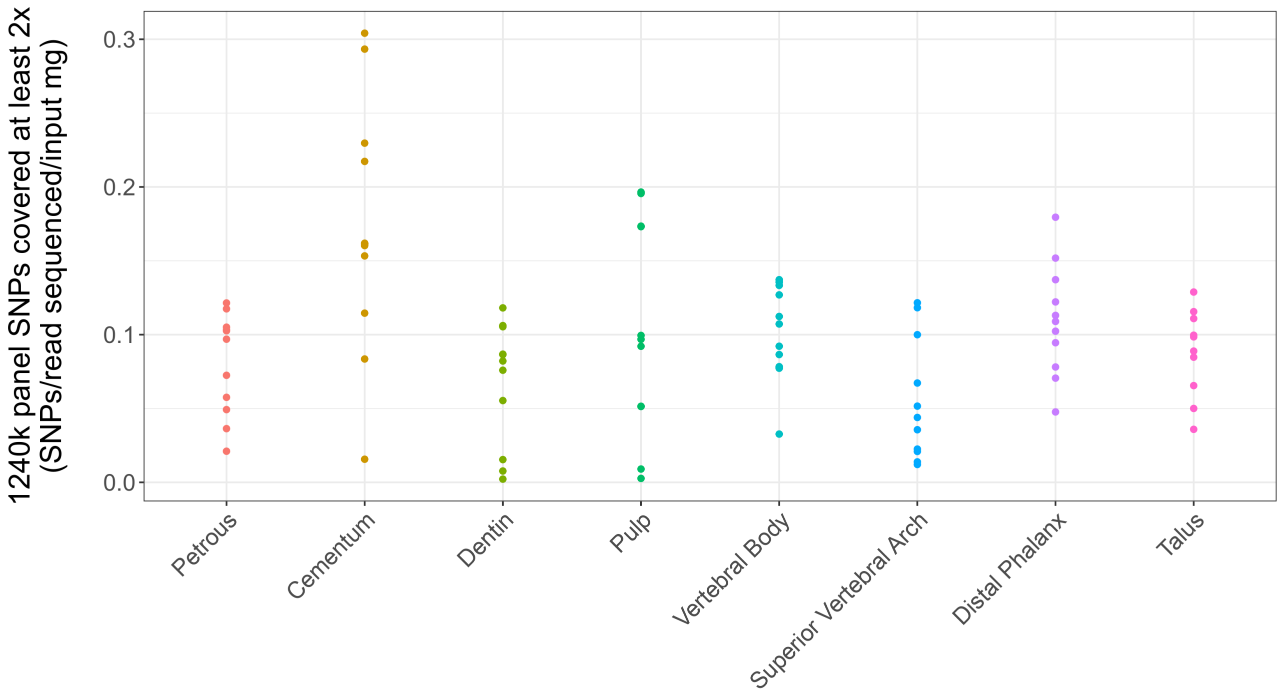


**Figure S16.** Number of 1240k panel SNPs covered at least 2x per read sequenced per mg of input material for all sampling locations, showing the increased richness of both the cementum and dental pulp chamber.

**2.5 Evaluation of metaphasic plates found in the two juvenile individuals included in the eleven individuals used for sampling.**

Sampling was also conducted on the metaphasic plates identified among the skeletal elements/individuals used for the main body of this study (from two individuals: in the femur, hip, and metacarpal of KRA004, in the femur and metacarpal of KRA005). As there were not enough instances of these features, they were excluded from the comparative analyses presented in the main body of this study. However, it should be noted that libraries stemming from metaphasic plates performed well in terms of proportion of human DNA recovered, reads mapping to the human genome per million reads, and estimated genomic coverage with respect to libraries from other sampling locations from the same element in that individual (as evidenced in the corresponding Supplementary File 1). As such, further study into aDNA preservation in skeletal features such as metaphasic plates in subadult individuals may be worthy of future consideration.
